# OPS-γδ: allogeneic opsonin-secreting γδT cell immunotherapy for solid tumours mediates direct and bystander immunity

**DOI:** 10.1101/2022.10.23.513387

**Authors:** D Fowler, M Barisa, A Southern, C Nattress, E Hawkins, E Vassalou, A Kanouta, J Counsell, E Rota, P Vlckova, B Draper, C Tape, K Chester, J Anderson, J Fisher

**Affiliations:** UCL Great Ormond Street Institute of Child Health, London, UK; UCL Cancer Institute, London, UK

## Abstract

T cell-based cancer immunotherapy has typically relied on membrane-bound cytotoxicity enhancers such as chimeric antigen receptors expressed in autologous αβT cells. These approaches are limited by tonic signalling of synthetic constructs and costs associated with manufacture of bespoke patient products. γδT cells are an emerging alternative chassis for cellular therapy, possessing innate anti-tumour activity, potent antibody-dependent cytotoxicity (ADCC) and minimal alloreactivity. We present an immunotherapeutic platform technology built around the Vγ9Vδ2 γδT cell chassis, harnessing specific characteristics of this cell type and offering an allo-compatible means of delivering cellular therapy that recruits bystander immunity. We engineered γδT cells to secrete synthetic opsonins and stabilized IL15 (stIL15). Using GD2 as a model antigen we show how opsonin-secreting Vγ9Vδ2 (OPS-γδ) have enhanced cytotoxicity and also confer this benefit on lymphoid and myeloid bystander cells. Reflecting the secreted nature of the engineered efficacy modules, the entire product rather than just the gene-modified fraction exhibited enhanced activation and cytotoxic profiles, superior persistence and proliferative capacity even upon repeated tumour challenge. Secretion of stIL15 abrogated the need for exogenous cytokine supplementation during expansion and further mediated functional licensing of bystander NK cells. Compared to unmodified γδT cells, stIL15-OPS-γδ cells exhibited superior *in-vivo* control of subcutaneous tumour and persistence in the blood. stIL15-OPS-γδ cells were further efficacious in 3D patient-derived osteosarcoma models, where efficacy could be boosted with the addition of immunomodulatory aminobisphosphonate drug, zoledronic acid. Together the data identify stIL15-OPS-γδ cells as a novel allogeneic platform combining direct cytolysis with bystander activation to effect solid tumour control.

**One Sentence Summary:** Armoured, opsonin-secreting OPS-γδ cell immunotherapy is built on the innate strengths of the Vγ9Vδ2 cell chassis for allogeneic solid tumour targeting.

## INTRODUCTION

Cellular immunotherapy using genetically modified T cells shows striking success against haematological malignancies*(1)*. Synthetic immunotherapeutic modules such as chimeric antigen receptors (CARs) linking tumour-associated antigen recognition to T cell effector function have been in development since the 1990s*(2)*. There has, however, been a relative paucity of clinical success against solid tumours, contributed to by the highly immunosuppressive tumour microenvironment and poor penetration of the tumor by engineered cells.

Vγ9Vδ2-γδT cells are a versatile chassis for cellular immunotherapy, possessing many helpful properties that include potent antibody-dependent cellular cytotoxicity (ADCC) capacity*(3–6)*, a tissue-tropic homing profile*(7)*, a range of innate tumour-sensing receptors minimizing the likelihood of tumour escape*(8)*. Furthermore, γδT cells are straightforward to expand to clinically useful numbers*(6, 9, 10)*, cause minimal graft-versus-host disease in the allogeneic setting*(11)* and, when present in the tumour microenvironment are a strong positive correlate with good clinical outcome *(12, 13)*.

Whilst progress has been made in “armoring” CAR-T with secreted cytokines, most T-cell engineering strategies rely on membrane-bound constructs. This confines any enhanced activity to the engineered cells; un-engineered cells in the product and bystander immune cells receive little if any benefit. We sought to overcome these challenges by designing an allo-compatible cell therapy platform that produces secreted mediators only. We demonstrate that the combination of engineered cytokine armouring and secreted opsonin production enhances both direct and bystander cytotoxicity, product persistence and phenotype.

## RESULTS

### γδT cells can be engineered to deliver multiple immune-active secreted payloads

For clarity, the nomenclature used to describe various effector cells in this manuscript is outlined in Table 1. Vγ9Vδ2 cells from healthy donors were expanded and transduced to express either a secreted scFv-Fc fusion protein (SFP) opsonin, a secreted IL15 construct, or both (Fig 1A); in all cases GFP was used as a transduction marker. We confirmed that opsonins targeting a range of tumor associated antigens (CEACAM5, CD20 and GD2) were detectable in transduced cell supernatant (Fig 1B & S1A), and chose to focus on the GD2-targetting binder 14G2a. GD2 is a proven immunotherapeutic target expressed on neuroblastoma, Ewing sarcoma, osteosarcoma and glioma *(14–19)*, with very limited expression in healthy neuronal tissue*(20–22)*. The restriction of GD2 expression to specific tissues facilitates use of isogenic GD2+/− lines for determination of antigen-specific effects. A mean 6.1±1.7 ng/mL anti-GD2 opsonin was detected in the supernatant of OPS-γδ cell culture at harvest (n=3 representative donors, determined by flow cytometry), equivalent to 3.97±1 ng/10^6^ cells (Fig S1B). OPS-γδ cells mediated enhanced antigen-specific cytotoxicity compared to unmodified γδ cells (Fig 1C & S1C). Whilst in overnight killing assays OPS-γδcells had lower cytotoxicity than matched 14G2a-28ζCAR-γδ cells, in longer term (7 day) challenge assays their ability to control SupT1-GD2 growth was equivalent to that of CAR-T (Fig S1D-E). OPS-γδ showed significantly lower expression of exhaustion associated markers TIM-3, LAG-3 and TIGIT than CAR-γδ at harvest of culture (Fig S2A and B).

**Table 1.** Nomenclature.

| *Nomenclature* | *Description* |
| --- | --- |
| OPS-$\gamma\delta$ cells | Vγ9Vδ2 cells expanded from zoledronic acid-stimulated PBMC and virally transduced to express opsonin-secretion constructs. Harvested on day 12 of expansion, unless otherwise specified. |
| stIL15-OPS-$\gamma\delta$ cells | Vγ9Vδ2 cells expanded from zoledronic acid-stimulated PBMC and virally transduced to express opsonin-secretion constructs as well as stIL15. Harvested on day 12 of expansion, unless otherwise specified. |
| Unmodified-$\gamma\delta$T cells | Vγ9Vδ2 cells expanded from zoledronic acid-stimulated PBMC, but not further modified. Harvested on day 12 of expansion, unless otherwise specified. |
| Freshly-isolated $\gamma\delta$T cells | Unstimulated Vγ9Vδ2 cells in freshly-isolated PBMC. |
| stIL15-14G2a-$\gamma\delta$ cells | stIL15-OPS-$\gamma\delta$ cells that express clone 14G2a SFP. |
| stIL15-isotype-$\gamma\delta$ cells | stIL15-OPS-$\gamma\delta$ cells that express an SFP of irrelevant specificity, acting as an isotype control for the secreted opsonin. |

**Figure 1.**
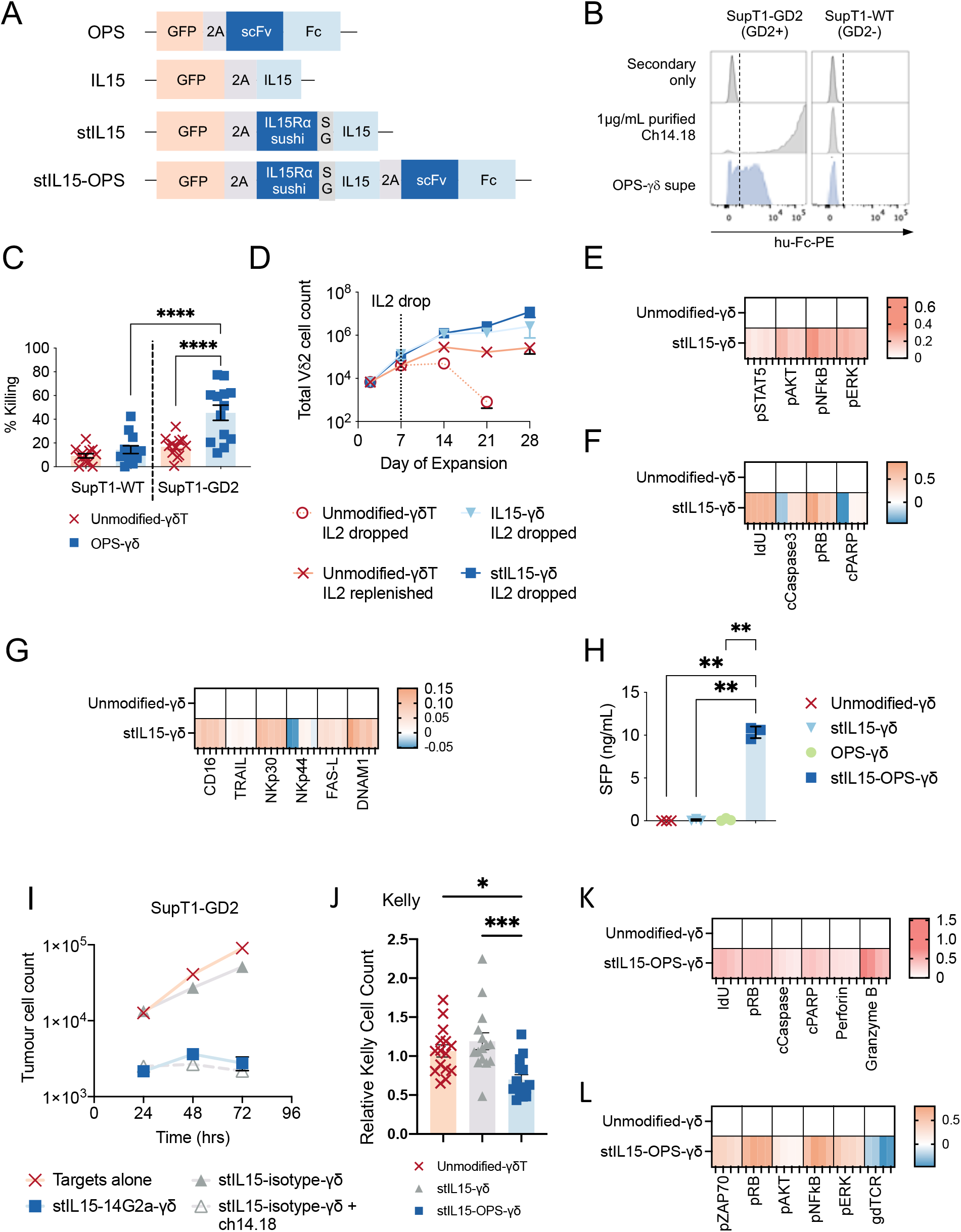
γδ T cells engineered to secrete opsonins and stabilized IL15 exhibit a favourable immunotherapeutic profile. (**A**) Bicistronic constructs encoding a GFP reporter gene in combination with SFPs, or tricistronic constructs encoding GFP, IL15 and SFP. (**B**) Flow cytometry of various antigen-positive tumor cells exposed to culture supernatant from either unmodified-γδ T or 14G2a OPS-γδ . Fluorochrome-conjugated anti-Fc or anti-IgG were used to detect SFP binding. 1ug/ml purified whole IgG1 against the relevant antigen was used as a positive control. Representative plots from 3-5 donors are shown. (**C)** Percentage killing of SupT1-WT and SupT1-GD2 cells by unmodified-γδ or 14G2a OPS-γδ (n = 13 across 8 donors) or 14G2a CAR-γδ (**G**, n=3 donors) in an overnight co-culture at an E:T of 1:1. **(D)**Vδ 2 cell counts over time during the expansion of unmodified-γδ T, IL 15-γδ and stIL15-γδ as measured by flow cytometry. Data are means±SEM for n=3 donors. (**E-G**) Expression of key markers of cell signaling (**E**), cell cycle and apoptosis (**F**) and cytotoxic function (**G**) commonly associated with IL15 signaling was assessed in unmodified-γδ T, IL15-γδ and stIL15-γδ on day 12 of expansion using mass cytometry. Differences are normalized to unmodified-γδ T as the baseline. Data shown in the heat maps EMD from n=5 donors. **(H)** Detection and quantification of 14G2a opsonin in supernatant from unmodified-γδ, 14G2a-γδ or stIL15-14G2a-γδ cultured for 72h was achieved using flow cytometry (Fig S1B for standard curve). **I**) Effect of adding either stIL15-14G2a-γδ, stIL15-isotype-γδ or stil15-isotype-γδ + 1 μg/ml Ch14/18 to growth of SupT1 -GD2. **(J)** Cytotoxicity assay comparing effects of unmodified γδ, stIL15 γδ or stIL15-14G2a-γδ on Kelly neuroblastoma cells. Relative tumor cell count was generated compared to tumor targets cultured alone. **(K-L)** Expression of key markers of proliferation, apoptosis and cytotoxic function was assessed in unmodified-γδ or stIL15-OPS-γδ on day 12 of expansion using mass cytometry. Differences are with reference to unmodified-γδ as the baseline, and heatmaps show EMD of n=4 across 2 donors.

IL15 is a commonly-used cytokine in γδT cell expansion protocols*(6, 9, 10, 23, 24)*, to which γδT cells become addicted*(25)*. Armoring of γδT cells with membrane bound IL15 is clinically beneficial*(26)* and the activity of IL15 can be enhanced by generating a fusion protein with the sushi domain of IL15Rα; we denote this stIL15. Such stabilized IL15 constructs have been used both as drugs in their own right and as armoring strategies for CAR-αβtherapies *(27–31)*. Transduction efficiencies were higher for the stIL15 construct compared to IL15 despite equal MOI of viral vector Fig S3A-B). Secretion of either IL15 or stIL15 rescued γδT cells from product viability and expansion collapse induced by the removal of IL-2 (Fig 1D & S3C), but stIL15 yielded much higher concentrations of detectable cytokine than native IL15 irrespective of IL-2 supplementation (Fig S3D). Due to the enhanced expansion dynamics observed in stIL15-γδ compared to IL15-γδ cells and the higher levels of detectable mitogen with stIL15, we progressed the stIL15 module for further testing.

We interrogated the effects of stIL15 on γδT cell signaling using mass cytometry followed by Earth Mover’s Distance analysis to determine differences in abundance of key signaling and phenotypic markers whilst remaining faithful to the single-cell resolution of the dataset*(32)*. Compared to unmodified-γδT cells, stIL15-γδ cells had upregulated pSTAT5, pAKT, pERK and pNFkB, consistent with IL15-driven responses (Fig 1E and Fig S4A). stIL15-γδ showed higher markers of proliferation (iododeoxyuridine (IdU) incorporation and pRB) without concurrent upregulation of apoptotic markers such as cleaved PARP or cleaved Caspase 3, suggesting that stIL15-γδ are more activated but no more prone to apoptosis than donor-matched unmodified γδ (Fig 1F and Fig S4B). Consistent with a more favorable tumor-targeting profile stIL15-γδ expressed more Fas-L, TRAIL, DNAM1 and NKp30, though levels of NKp44 were not significantly affected (Fig 1G and Fig S4C).

Having separately demonstrated the benefit of OPS- and stIL15-modules, we combined them to produce stIL15-OPS-γδ. Transduction efficiency remained high (75±2.5% mean±SEM for n=13 over 8 representative donors) and co-expression of stIL15 and opsonin was demonstrated by ELISA and flow cytometry, respectively (Fig S4D, E). When stIL15-OPS-γδ were re-seeded into fresh media after harvest of expansion 12 days post-initiation, opsonin yield 72h later was substantially higher than that observed from OPS-γδ in the same conditions (Fig 1H), with a yield of 1.55±0.05ng/10^6^ cells seeded (mean of n=3 representative donors). This suggests that whilst OPS-γδ produce opsonin initially, production falters, whereas stIL15-OPS-γδ sustain opsonin secretion. stIL15-OPS-γδ showed enhanced expansion and Vδ 2 purity compared to unmodified γδ T cells (Fig S4F-G). Cytotoxicity of stIL15-14G2a-γδ was both antigen-specific and potent (Fig 1I) and was not confined to the isogenic model system; showing enhanced killing of GD2+ Kelly (neuroblastoma) which was not observed with stIL15-γδ cells (Fig 1J).

Compared to unmodified-γδ T cells, stIL15-OPS-γδ retained the increased IdU incorporation and pRB observed in stIL15-γδ but also had increased levels of apoptosis-associated cleaved caspase 3 and cleaved PARP (Fig 1F). Consistent with their enhanced activation and cytotoxicity, stIL15-OPS-γδ had higher levels of Perforin and Granzyme B compared to unmodified γδ (Fig 1K and Fig S5A, C). Indicative of activation-related transcriptional changes, stIL15-OPS-γδ had elevated pNFkB levels, also showing intracellular signaling typical of T cell activation, including ITAM-dependent pZAP70 and pAKT, as well as pERK with relative downregulation of the γδ -TCR (Fig 1L and Fig S5B).

To interrogate the autonomous growth potential of stIL15-14G2a-γδ cells, we cultured the cells target-free for 28 days. To avoid under-representing autonomous expansion due to culture overgrowth, cells were split and supplemented with fresh cytokine-free complete media as required. Most of the proliferation took place in the first two weeks of culture, followed by a plateau in expansion, indicating that leukemic proliferation had not been induced (Fig 2A). Cytotoxicity of stIL15-14G2a-γδ was sustained over time in a 4-day re-challenge assay (Fig 2B,C). Following clearance of SupT1-GD2, stIL15-14G2a-γδ effectively controlled a re-challenge with SupT1-GD2 but not SupT1-WT (Fig 2C). Having established tumor control, stIL15-14G2a-γδ cells had a lower growth rate in the presence of tumor cells than target-free conditions, but nonetheless continued to proliferate in the face of challenge with tumour (Fig 2D). Unmodified γδ T did not expand in either condition (Fig S6A). We, therefore, concluded that stIL15-14G2a-γδ cells maintain tumor control through sustained cytotoxic function whilst retaining capacity to persist and even expand.

**Figure 2.**
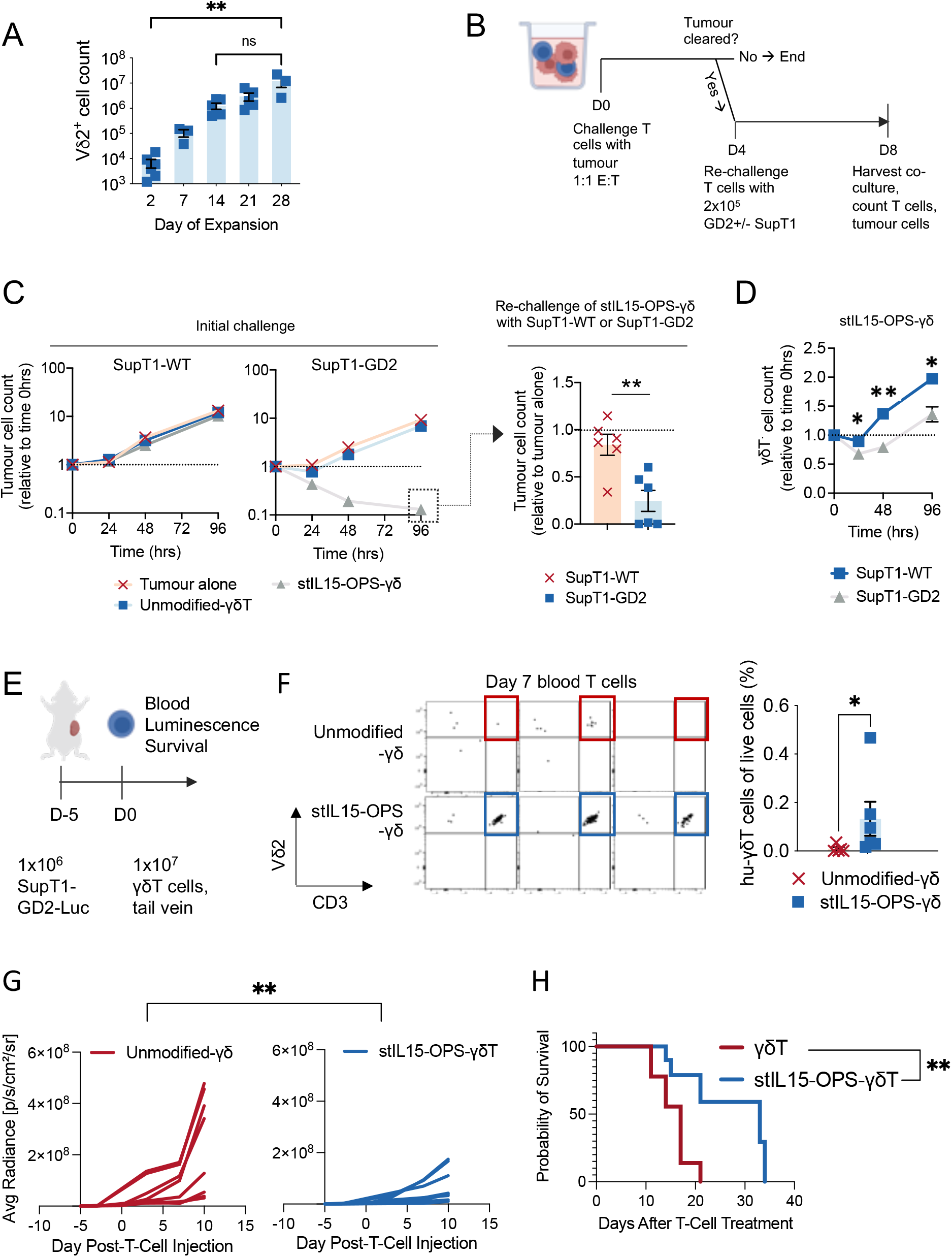
A single IV dose of stIL15-OPS-γδ cells is efficacious against a model of subcutaneous GD2-positive tumour. (**A**-**D**) SupT1-GD2 cell counts over time in 1:1 co-cultures with stIL15-OPS-γδ secreting either 14G2a or isotype SFP—designated stIL15-14G2a-γδ or stIL15-isotype-γδ, respectively—as measured by flow cytometry. 1μg/ml ch14.18 was added to some conditions. Data in (**A**) shows means of experimental triplicates from one representative donor. (**B**) Schematic depicting a re-challenge experiment to test the serial killing capacity of stIL15-OPS-γδ . (**C**) SupT1-WT and SupT1-GD2 cell counts during initial 1:1 E:T challenge and then following post-clearance re-challenge; n=3 for tumor alone, n=6 across 2 donors for co-cultures, comparison by unpaired t-test). (**D**) Vd2+ cell counts in stIL15-OPS-γδ preparations in presence of either SupT1-WT or SupT1-GD2 (mean ± SEM of n=6 across 2 donors, comparison by 2-way ANOVA). CD3^+^Vδ 2^+^ cell counts over time in target-free expansion cultures. Individual data points and means±SEM for n=3-6 independent donors are shown, comparison by Kruskall-Wallace test with Dunn’s multiple comparison correction. (**E**) NSG mice were challenged with 1×10^6^ luciferase-expressing SupT1-GD2 cells (SupT1-GD2-Luc) in Matrigel. Five days later, mice were treated with 1×10^7^ unmodified-γδT or 14G2a SFP-secreting stIL15-OPS-γδ . Blood samples were taken to track cell survival, and serial bioluminescence measurements used to track tumor growth and efficacy over time. (**F**) Flow cytometry was used to detect human CD3^+^Vδ 2^+^γδ T cells in day 7 blood samples. Representative plots from 3/6 mice per group are shown along with individual data points and means±SEM for all mice. **(G)** Bioluminescence measurements plotted as average radiance over time for individual mice. Data is pooled from two independent experiments: a pilot study containing three mice per group and a follow-up study containing 6 mice per group. (**H**) When tumors exceeded 1.5cm by vernier caliper measurement in any one direction, mice were culled, and this data used to calculate mock survival curves (n=8) per group.

To test stIL15-OPS-γδ *in-vivo* efficacy in the absence of exogenous cytokine, NOD.Cg-*Prkdc^scid^ Il2rg^tm1Wjl^/SzJ* (NSG) mice bearing subcutaneous SupT1-GD2 tumors received a single dose of 1×10^7^ intravenous unmodified- or stIL15-OPS-γδ cells without further cytokine support (Fig 2E). Tumor progression was monitored using bioluminescence and caliper measurements. Blood samples collected on day 7 post-T cell administration showed more Vδ 2-γδ T cells in the blood of stIL15-OPS-γδ -treated mice than unmodified-γδ treated mice (Fig 2F). Tumor luminescence and size comparisons were censored after day 10 following humane cull of animals with large tumors. At day 10, tumor luminescence and volume were lower in animals treated with stIL15-OPS-γδ compared to unmodified-γδ T cells (Fig 2G, S6B-D), and stIL15-OPS-γδ treatment prolonged animal survival over treatment with unmodified-γδ T cells (Fig 2H).

### Secreted transgene products relieve dependence on transgene expression for cellular enhancement

Evaluation of the differential effects of membrane-bound and secreted constructs on T cell signaling requires a means of accurately determining the dependence of signaling markers on the presence of transgenes at a single-cell level. As representative comparators we used two 2^nd^ generation CAR-T products, targeting either CD33 or ALK, with human-Fc stalks, CD28 transmembrane regions (detectable using anti-human Fc antibodies) and CD28-CD3ζendodomains. For CAR-T we analyzed the differences in marker abundance by comparing CAR+/− cells in the same sample, whereas stIL15-OPS-γδ were compared to donor and time-matched unmodified-γδ cells to eliminate the confounding effect of secreted product.

Simple correlative measures overemphasize the influence of denser areas of a distribution, neglecting biologically informative outliers. We therefore used Density rescaled visualization (DREVI) plots and Density Rescaled Estimates of Mutual Information (DREMI) scores *(33)* to illustrate the relationship between transgene expression and signaling markers. DREMI describes the strength of relationship or “edge strength” between two markers. A brief overview of DREVI and DREMI derivation is shown in Fig 3A, and the sample handling for these comparisons is shown in Fig S7. A high DREMI score for a given edge X→M indicates that M is highly dependent on X, whereas a low DREMI score indicates that they are independent of each other.

**Figure 3.**
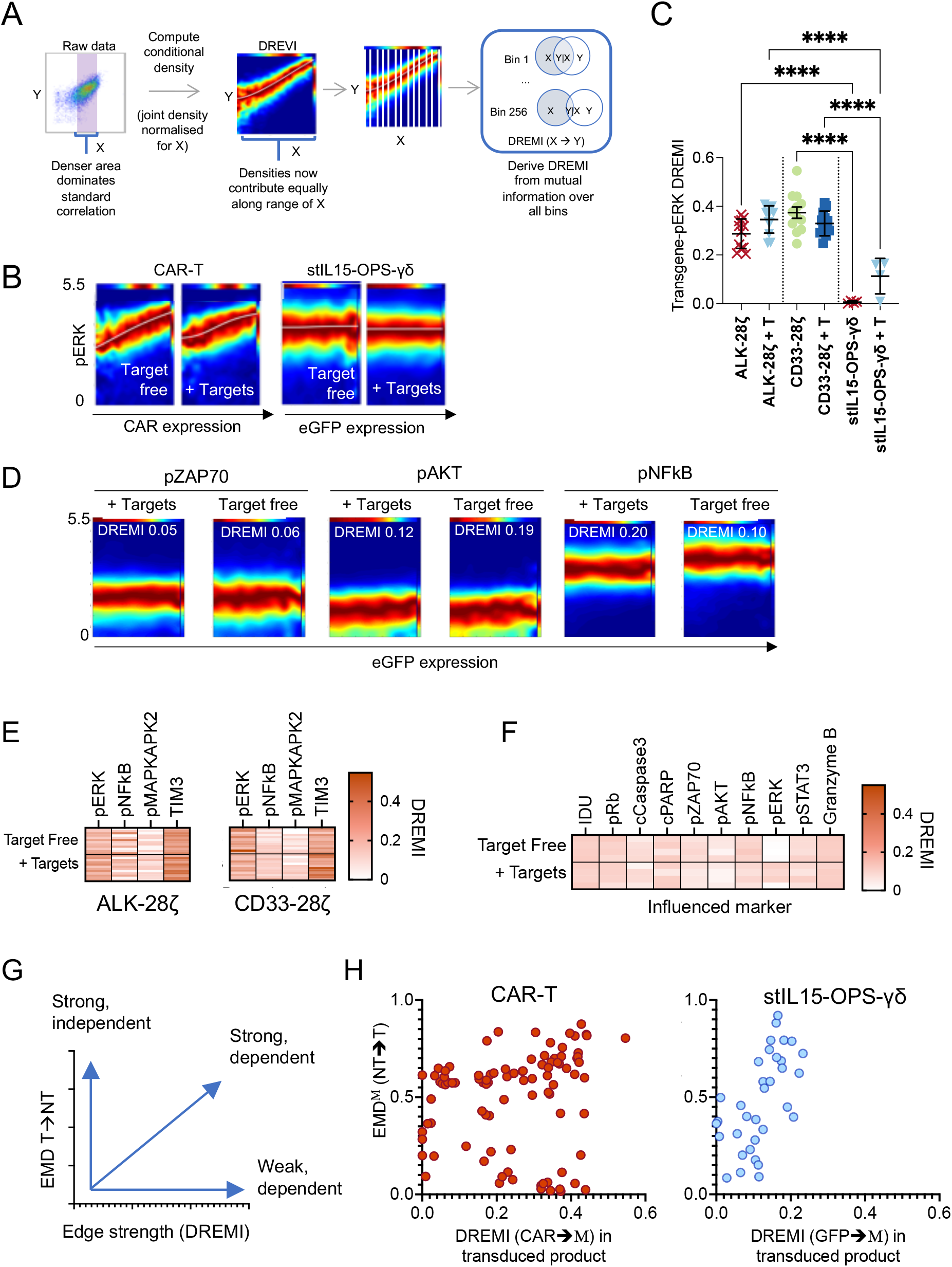
Secreted transgene products relieve dependence on transgene expression for cellular enhancement. (**A**) Example DREVI/DREMI analysis of correlation between molecules X and Y. A high DREMI score with a steep gradient on DREVI indicates that Y expression is highly dependent on X. **(B)** Comparison of X-pERK DREVI plots and DREMI scores for representative CAR-T cells (expressing ALK-28ζ CAR or CD33-28ζ CAR, n=12 across 3 donors) and stIL15-14G2a-γδ (n=4 across 2 donors) in the presence of absence of targets expressing the relevant antigen, where X is the marker of transduction. The first component of the DREMI score and X-axis of the DREVI plots is either directly stained CAR expression, or GFP in the case of stIL15-OPS-γδ which lack a membrane-bound marker. **(D)**s tIL15-OPS-γδ pERK is much less dependent on construct expression (comparison by one-way ANOVA with Tukey’s correction); this lack of dependence is replicated across other canonical signalling nodes. **(E-F)** DREMI scores describing the influence of transgene expression on markers known to change expression in response to transgene activity in CAR-T (E) and stIL15-OPS-γδ (F). **(G)** Across all markers measured, plotting the size of the effect (as measured by EMD) against the dependence of the effect on transgene expression (as measured by DREMI) gives information on strength and transgene dependence. **(H)** CAR-T signalling effects are highly dependent on CAR expression at a single-cell level, whereas stIL15-OPS-γδ lead to strong signals which are not dependent on transgene expression.

DREVI was used to visualize the relationship between pERK and transgene expression; *(12)* GFP was used as a marker of stIL15-OPS expression whereas membrane-bound CARs were directly stained. CARàpERK DREVI plots exhibited a clear increase in pERK as CAR expression increased, whereas stIL15-OPS-γδ showed steady levels of pERK throughout all cells in the product (Fig 3B). These differences were reflected in the DREMI scores, CAR-pERK DREMI scores were high and showed little plasticity, whereas GFP-pERK scores were low (Fig 3C). Similar DREVI profiles were observed in stIL15-OPS-γδ for GFPàpZAP70, GFPàpAKT and GFPàpNFkB (Fig 3D). Focusing on species known to be influenced by 28ζ-CAR signaling, we observed high DREMI scores for CARàpERK, CARàpNFkB and CARàTIM-3 edges for both CAR-T products analyzed, in particular CARàpERK and CARàTIM-3 (Fig 3E). In contrast, despite evidence that stIL15-OPS-γδ engineering increases abundance of many signaling species (Fig 1K-L), the strength of relationship between these markers and GFP expression remained low (Fig 3F).

Taken together, the “effect of transduction”, which can be derived from the change in marker abundance (EMD) between transduced and unmodified cells, and the “dependence on transgene” which can be derived from the DREMI score for edges beginning with the transduction marker (CARàA, B, C or GFPàA, B, C) give an overall picture of signaling behavior in these two engineered product types. Markers with high transduction induced EMD and high DREMI scores indicate strong signals only in cells expressing the transgene (Fig 3G).

Signals in unstimulated ALK-28ζ and CD33-28ζCAR-T and stIL15-OPS-γδ products were analyzed. The transduction-induced EMD for a given marker (M) was plotted against that marker’s dependence on the transgene (DREMI transgeneàM, using all markers where an effect of transduction had been identified. The CAR-T plot was enriched in the “strong dependent” area, indicating that the tonic signaling observed in the CAR-T population was highly dependent on the presence of the detectable CAR molecules in the cell. Conversely, whilst there were strong signals detected in stIL15-OPS-γδ, the plots were enriched in the “strong, independent” region, demonstrating that the detected changes did not require the transgene to be present in an individual cell for that cell to benefit (Fig 3H)

Having demonstrated that stIL15-OPS-γδ confer benefits to non-transduced cells within the product, we investigated if these benefits could be conferred to other bystander immune cells (Fig 4A). We evaluated the ability of OPS-γδ and stIL15-OPS-γδ supernatant to engage the antigen-specific cytotoxicity of a range of ADCC-competent cells, including unmodified γδ T cells, NK cells and macrophages (Fig 4B & Fig S8A). To provide a secondary line of evidence for the antigen-specificity of bystander activity enhancement we used the stIL15-isotype-OPS-γδ system first described in Fig 1I. stIL15-14G2a-γδ supernatant mediated potent killing of antigen-positive targets by stIL15-isotype-γδ (Fig 4C), demonstrating that the antigen-specific responsiveness was not impeded by production of an irrelevant opsonin. Supernatant from 1^st^ generation 14G2a OPS-γδ cells (without stIL15) induced antigen-specific cytotoxicity in unmodified γδ T cells (Fig S8B), and neutrophils (Fig S8C); but did not enhance NK cell cytotoxicity - despite their demonstrable ADCC activity mediated by the addition of 1 μg/mL of ch14.18 mAb (Fig S8D). IL15 has been described as a potent licenser of NK cell effector function and differentiation*(34)*; stIL15-OPS-γδ supernatant mediated substantial antigen-specific NK cell cytotoxic enhancement (Fig 4C). Supernatant from stIL15-OPS-γδ but not stIL15-γδ further mediated efficient clearing of GD2-positive tumor cells by macrophages, while GD2-negative targets were spared, highlighting a requirement of both opsonin and target antigen (Fig 4E).

**Figure 4.**
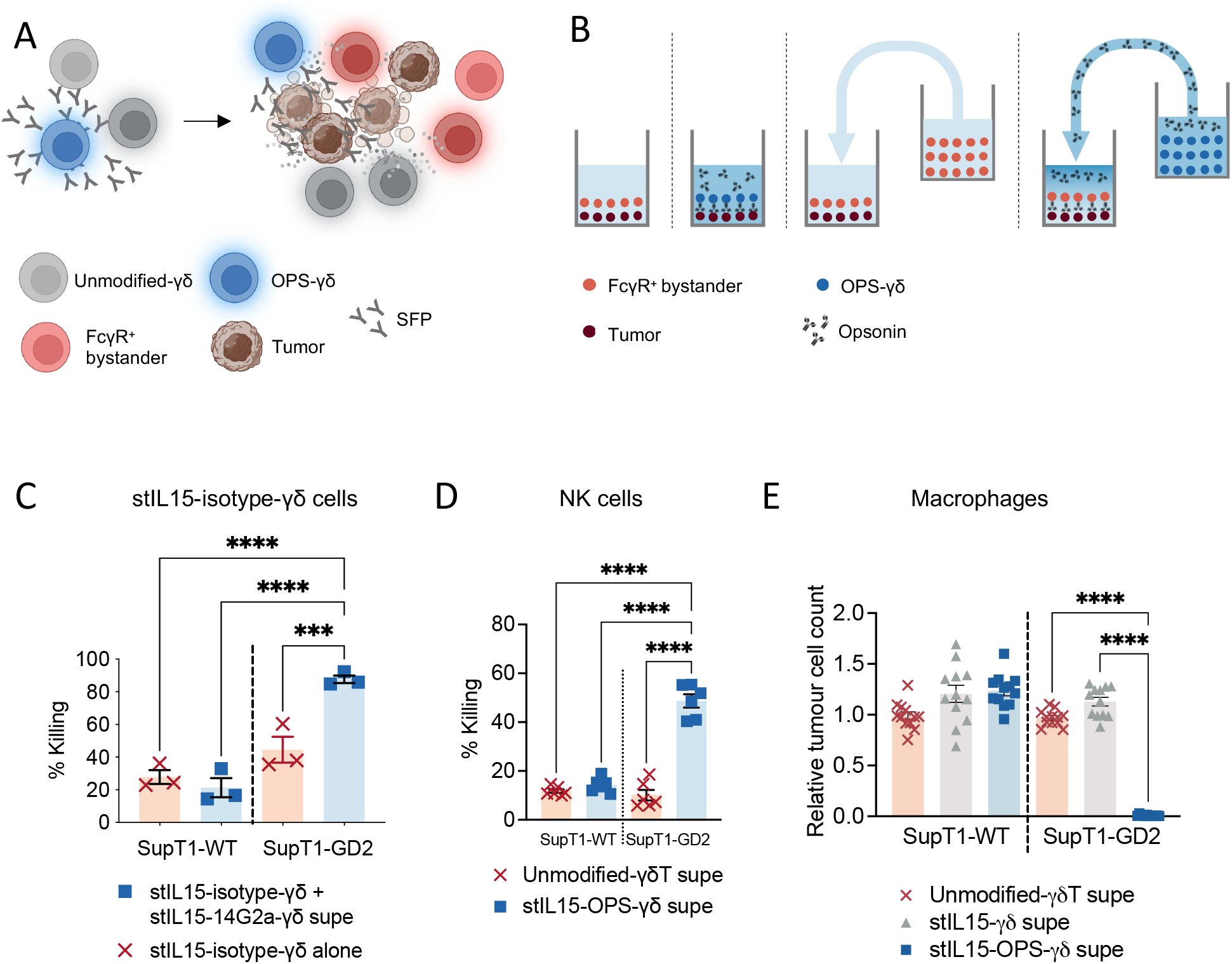
stIL15-OPS-γδ recruit bystander cells to elicit ADCC and/or ADCP responses against antigen-positive tumor target cells. **(A)** Schematic depicting the bystander recruitment potential of stIL15-OPS-γδ . **(B)** Schematic showing experimental setup for determining the effect of secreted product **(C-E)** Flow cytometry was used to measure the effect of supernatant from unmodified γδ T or stIL15-OPS-γδ on percentage killing of SupT1-WT and SupT1-GD2 cells co-cultured with various effectors including of resting NK cells (**C**, n=6 supernatants from 3 donors), unmodified expanded γδ T (**C**, n=9 across 5 donors), **(D)** Stem cell-derived neutrophils (mean ± SEM of n=3 supernatants from 3 donors) and **(E)** resting NK cells (n=9 supernatants from 3 donors or n=9 for ch14.18 control), stIL15-isotype-Vδ 2 (D, n = 3 supernatants from one donor monocyte-derived macrophages (**H**, n=12 supernatants from 4 donors). Individual data points are shown as well as means±SEM, all comparisons are by one-way ANOVA with Tukey’s multiple comparison correction.

### stIL15-OPS-γδ phenotype favors ADCC and osteosarcoma-homing

After initial expansion with zoledronic acid and IL2 but before transduction, Vδ 2-γδ T cells expressed high levels of CD16b and CD64 FcγR’s, with moderate CD32 expression. CD16a was only present on a very small subset of cells (Fig 5A). Following engineering to form OPS-γδ, CD16b expression was substantially decreased, with a marked but variable increase in CD16a expression, such that this isoform comprised the entirety of CD16 expression by D10 of culture. Notable changes did not present for CD32 and CD64 expression (Fig 5A).

**Figure 5.**
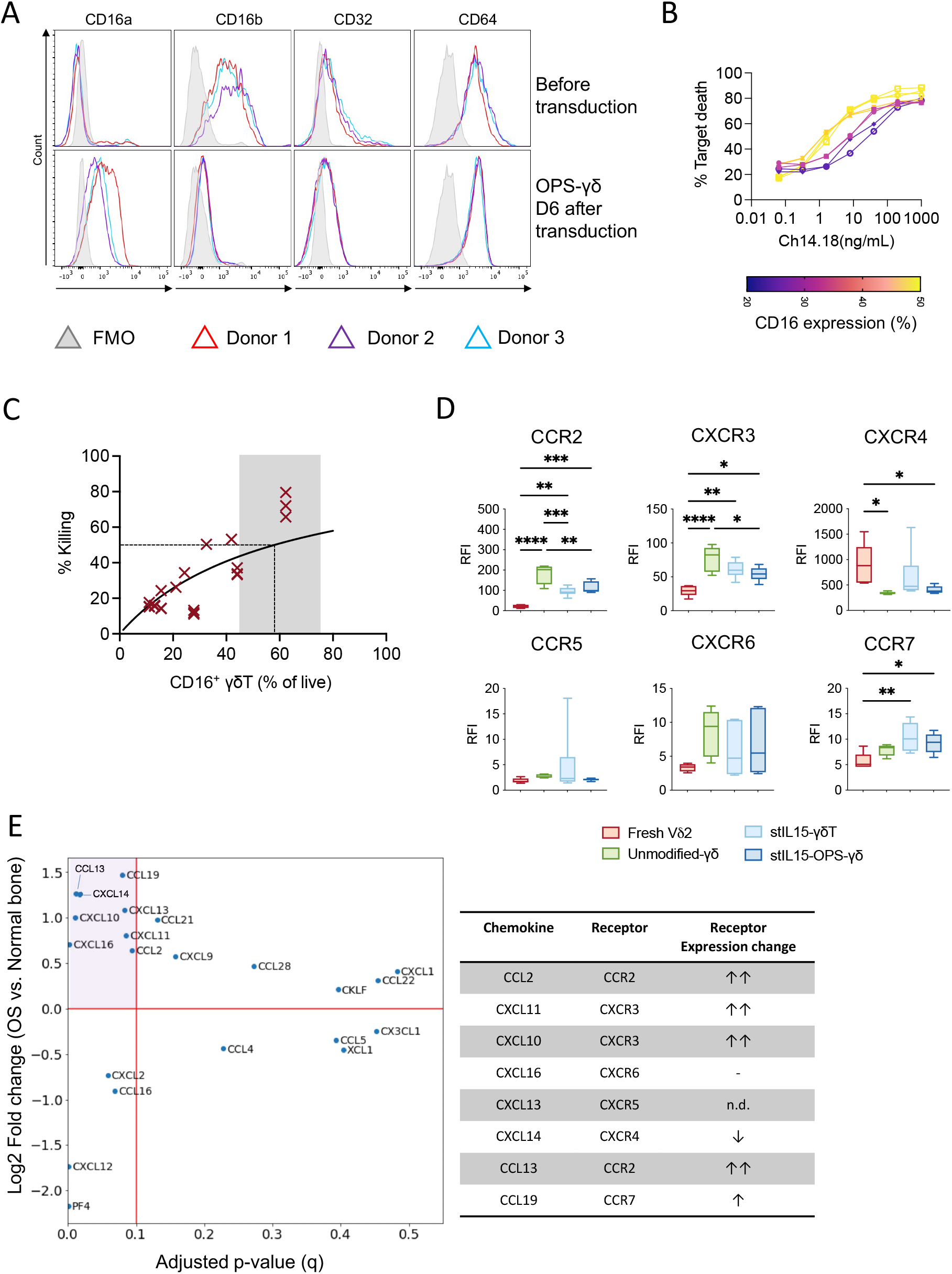
stIL15-OPS-γδ phenotype favors ADCC and osteosarcoma-homing. **(A)** Expression of CD16a, CD16b, CD32 and CD64 on Vδ 2-γδ T cells after stimulation with 5μM zoledronic acid + IL2, before and 6 days after transduction to form OPS-γδ . **(B)** Killing of SupT1-GD2 by unmodified γδ in the presence of varying concentrations of Ch14.18 (ranging from 0.064 ng/ml to 1μg/mL). n=8 across 4 donors, donor lines are colored by the expression level of FcγRIII (CD16) on Vδ 2 in the starting PBMC population. **(C)** Killing of SupT1-GD2 by unmodified γδ in the presence of 1.6ng/ml Ch14.18, plotted against levels of FcγRIII (CD16) on the corresponding donor’s Vδ 2 after expansion. Least squares curve fitting was used to determine the EC^50^ for CD16 expression, n=22 across 10 donors. **(E)** CXCR3, CCR7, CCR5, CCR2, CXCR6 and CXCR4 expression on gated Vδ 2^+^ cells within fresh PBMC vs either unmodified-γδ, stIL15-γδ or stIL15-OPS-γδ on day 12 of expansion as assessed by flow cytometry. Box and whisker plots show mean and 5-95^th^ centile for n=6 donors; table indicates the receptor expression changes in the context of osteosarcoma chemokine enrichment. **(D)** Differential expression of chemokines in osteosarcoma compared to normal bone from the same patient (n=18)

Despite a consistent transduction efficiency (56.5±4.8%, n=18 across 10 donors), cytotoxicity of OPS-γδ cells varied (Fig 1C). Because CD16a expression showed donor-donor variability whereas CD32 and CD64 expression was consistent, we hypothesized that variations in the interplay between low opsonin concentration and CD16 expression may govern Vδ 2 ADCC and thus OPS-γδ cytotoxicity. γδ T cells with higher CD16 expression mediated substantially higher cytotoxicity at low, physiologically-achievable and OPS-γδ -relevant opsonin concentrations in the 1-10ng/mL range (Fig 5B). At 1.6 ng/mL Ch14.18, the percentage of CD16 expression required to achieve 50% target killing was 58.1% (95% CI 44.9-76.6%) with strong correlation between cytotoxicity and CD16 expression (Pearson R^2^=0.77, p<0.0001, Fig 5C, FcγRIII gating shown in FigS9A). CD16 expression is, therefore, a potentially useful biomarker for selecting optimal allogeneic cell therapy donors for OPS-γδ product manufacture, and subsequent cytotoxicity data is presented on CD16^hi^ (>50% expression) γδ T cell donors. This correlation was lost at high (1μg/mL) Ch14.18 mAb concentration, where the percentage CD16 expression needed to mediate 50% killing at 1:1 E:T ratio was a low 6.4% (95% CI 4.5-8.8%), to the extent that the curve rapidly plateaued with poor correlation (Pearson R^2^=0.2, p =0.0094, Fig S9B). Transduction with GFP-IL15, GFP-stIL15 or GFP-stIL15-14G2a had minimal effect on CD16 expression compared to unmodified-γδ controls (Fig S9C). γδ T cell ADCC capacity has also been linked to their differentiation state*(3)*, but transduction with either eGFP-stIL15 or eGFP-stIL15-14G2a had no effect on CD45RA or CD27 expression compared to unmodified γδ, indicating no additional pressure towards terminal differentiation (Fig S9D).

Both stIL15-γδ and stIL15-OPS-γδ retained an activated γδ T phenotype consistent with enhanced tumor homing and chemotaxis compared to freshly-isolated peripheral Vγ9Vδ 2 cells. Both increased expression of chemokine receptors CCR2 and CXCR3 (Fig 5D). stIL15-OPS-γδ had increased CCR7 expression, potentially indicative of lymph node homing that may be consistent with the acquisition of a dendritic cell-type antigen presenting capacity of Vγ9Vδ 2 cells that has been previously described for both modified and unmodified activated Vγ9Vδ 2 cells*(7, 35–39)*. stIL15-OPS-γδ downregulated CXCR4, a receptor typically associated with tissue homing including to the bone marrow*(40)*, suggesting an enhanced tendency to remain in the periphery. Chemokine expression profiles in transcriptomic data from 18 patient osteosarcoma samples were compared to patient-matched normal bone using DeSeq2 implemented in Python (data from *(41)*, GEO accession GSE99671); 8 chemokines showed enhanced expression in osteosarcoma. 5 of these chemokines matched with receptors which had enhanced expression in stIL15-OPS-γδ (CCL2, CXCL10, CXCL11, CCL13 and CCL19, Fig 5E).

### Efficacy, mechanism and translational scalability of armored OPS-γδ against primary osteosarcoma

To be translatable to clinic, stIL15-OPS-γδ manufacture must be scalable and show activity against primary tumor exhibiting variable target antigen expression. Vγ9Vδ 2 cell expansion, transducibility and viability were evaluated in scalable GMP-compatible G-Rex® vessels (Fig S10A). High product viability (>80%), purity (>90% Vδ 2 cells of live cells) and transduction efficiency (>60%) were maintained in this system. Plated at 2×10^6^ PBMC/cm^2^, the system yielded 11.7±2.8×10^6^ γδ T cells cells/cm^2^ during a 12 day manufacturing process, corresponding to a total yield of ~5.6×10^9^ from a G-Rex®500M. these cells retained high cytotoxic functional capacity against GD2-positive SupT1 targets (Fig S10B). Three patient-derived OS lines *(42)* (kind gift from Dr Katia Scotlandi, Instituto Ortopedico Rizzoli, Bologna, Italy) were assessed for their levels of GD2 expression by flow cytometry (Fig 6A), demonstrating variable and heterogenous expression similar to that which has been described clinically*(43)*. We evaluated the mechanism and efficacy of stIL15-OPS-γδ responses to osteosarcoma using a combination of 2D and 3D co-culture systems. Modelling of tumor/immune cell interactions in a 3D culture system provides an opportunity for more biomimetic culture that can be analyzed at single-cell resolution using CyTOF (Fig 6B) *(53, 54)*.

**Figure 6.**
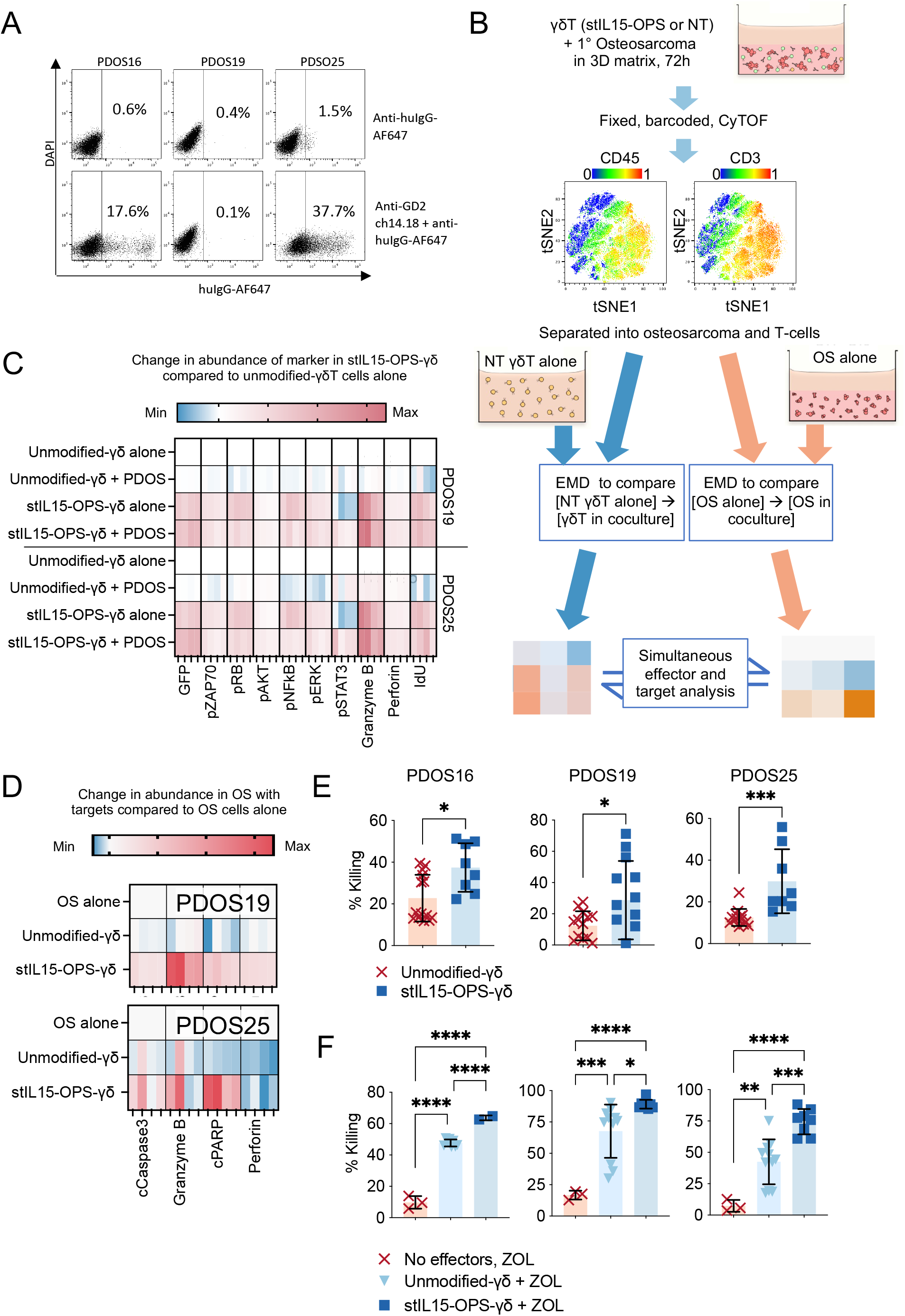
Armored OPS-γδ show activity against primary osteosarcoma that can be boosted using osteotropic compounds. (**A**) Expression of GD2 on 3 patient derived osteosarcoma lines. **(B)** Schematic of experimental setup for OS 3D co-culture. γδ T-cell preparations were confirmed to be >99% CD3+ and >90% Vδ 2+ prior to co-culture. Following CyTOF analysis, T-cells and OS were separated using CD45 and CD3, with OS being CD3-/CD45-. EMD was used to explore differences in abundance of key markers, compared to matched donor control samples cultured alone but in the same media conditions. (**C**) EMD analysis of signaling and functional markers in stIL15-OPS-γδ or unmodified γδ in the presence or absence of OS, using donor-matched unmodified γδ cultured alone as the baseline. **(D)** EMD analysis of cytotoxicity and apoptotic markers in OS in the presence or absence of either unmodified γδ or stIL15-OPS-γδ, using OS monoculture as the baseline. Data shown in all heatmaps is n=4 over 2 donors, with further data and statistical analyses in Fig S11A-B. **(E)** Cytotoxicity of unmodified γδ (n=14 across 6 donors) or stIL15-OPS-γδ (n = 8 across 4 donors) against PDOS lines was determined using flow cytometry. **(F)** Following overnight pre-treatment with 5μM zoledronic acid, cytotoxicity of unmodified γδ (n=11 across 6 donors) or stIL15-OPS-γδ (n = 8 across 4 donors) against PDOS lines was determined using flow cytometry, background death following zoledronic acid treatment (n = 3) is also shown.

Baseline enhanced phosphorylation of key activatory signaling nodes including ZAP70, AKT, and ERK in stIL15-OPS-γδ cells was preserved in the presence of both osteosarcoma lines, with additional significant increases in pAKT and reduction in pNFkB in response to GD2-expressing PDOS25 but not GD2-ve PDOS19 (Fig 6C, Fig S11A,B). Most target-induced effects were however GD2-autonomous, perhaps reflective of intermediate GD2 expression on PDX25 and the multifactorial innate reactivity of stIL15-supported γδ T cells against tumor. Expression of perforin was higher in stIL15-OPS-γδ cells in the presence of OS, and these target-induced increases were minimal or absent in unmodified-γδ T cells. Whilst co-culture with OS slightly reduced levels of pRb, levels were still significantly higher than in unmodified-γδ T cells in presence or absence of tumor. Engineering with stIL15-14G2a was associated with reduced pSTAT3 in the absence of targets, a pattern which was reversed in target presence (Fig 6C). Enhanced STAT3 signaling has previously been reported to confer resistance to activation-induced cell death in γδ T cells*(44, 45)*, supporting their proliferation; such effects would be consistent with the observed ability of stIL15-14G2a-γδ to persist and proliferate after repeatedly clearing tumor challenge (Fig 2C).

EMD analysis of osteosarcoma in co-culture was performed relative to matched OS monocultures. Relative to tumor alone and co-culture with unmodified-γδ, stIL15-OPS-γδ induced substantial accumulation of granzyme B in PDOS19 and upregulated cleaved PARP in both OS 3D cultures (Fig 6D). To contextualize this mechanistic information, we evaluated stIL15-OPS-γδ cytotoxicity against PDOS; stIL15-OPS-γδ cells outperformed unmodified-γδ T cells at killing OS. The enhancement was more consistent and reached greater significance when GD2 was present on the targets (Fig 6E). *(58, 59)*. This innate responsiveness could be boosted further with 5μM zoledronic acid pre-treatment, which was highly effective at sensitizing OS to killing by unmodified-γδ T cells. Maximal levels of cytotoxicity were achieved upon co-treatment with zoledronic acid and stIL15-OPS-γδ cells (Fig 6F).

## DISCUSSION

A major constraint of classical cell immunotherapies is the restriction of engineered moiety advantage to only the transduced portion of the medicinal product. By relying on membrane-bound moieties such as CARs or TCRs, conventional T-cell immunotherapies always contain an inactive fraction of cells, often up to 50%. Cytokine armoring is a step towards liberation but does not address the fundamentally membrane-bound nature of most “cytotoxicity switches”. The OPS-γδ therapeutic platform was designed to perform two tasks: (i) to effectively kill tumor targets directly, and (ii) to deliver an immunologically active payload to the tumor microenvironment, engaging cytotoxicity of un-engineered bystanders.

Autologous immunotherapies are associated with high cost, strained logistics and clinical challenges during the manufacturing period. Allogeneic, “off-the-shelf” therapies are an area of active development, and recent data has demonstrated the safety of allogeneic γδ T cells as a cancer therapeutic*(11, 46)*. Whilst approaches such as CAR-γδ are gaining traction, simply transferring technology designed for an αβT cell chassis to a γδ T cell neglects their innate properties and fails to capitalize on their innate tumor-reactivity and ADCC competence.

Here, we demonstrate a “chassis-faithful” engineering strategy for γδ T cells which not only enhances the properties of the engineered cells but also confers benefit on un-engineered cells in the product and on bystander immune cells from other lineages. The engineering strategy overcomes key limitations in γδ T cell product manufacture such as addiction to exogenous cytokines and does not confer the exhausting tonic signals seen with conventional CAR-T approaches*(47–49)*.

We demonstrate product behavior in the context of an isogenic model system and in the context of patient-derived osteosarcoma. Osteosarcoma is the commonest primary bone tumor in adolescents; current therapies involve highly toxic chemotherapy with life-altering surgery and patients frequently relapse with metastatic disease. The tropism of Vγ9Vδ 2 cell efficacy-enhancing compounds such as zoledronic acid for hydroxyapatite (a key component of bone) forms a rationale for using Vγ9Vδ 2-γδ T cell-based therapies in the context of such bone-forming tumors.

γδ T could be engineered to secrete two immunologically active species without leading to excessive cellular stress. Secretion of two separate moieties led to greater cellular stress than secretion of one – as evidenced by increased cleaved PARP and cleaved caspase in stIL15-OPS-γδ, not observed in stIL15-γδ . The proliferative benefits of IL15 armoring were not abrogated in the face of this however, IdU incorporation remained high as did proliferative capacity. Furthermore stIL15-OPS-γδ were able to continue proliferating whilst also controlling tumor growth, demonstrating a resistance to activation-induced cell death. The addition of stIL15 armoring led to a substantial enhancement in opsonin production by stIL15-OPS-γδ compared to OPS-γδ, most likely due to the enhanced cellular health imparted by stIL15 armoring.

Due to the marked inter-species differences between human and murine γδ T cells, modelling efficacy and bystander engagement by stIL15-OPS-γδ in immunocompetent animals is not possible. We were however able to demonstrate the enhanced survival of stIL15-OPS-γδ in an unsupported environment. Murine models of human γδ T cell therapy typically include dosing with cytokines such as IL2 or γδ T stimulants such as zoledronic acid. In the absence of any such support, a single dose of stIL15-OPS-γδ was able to persist and confer enhanced survival and control of highly aggressive tumors. In such an immunodeficient model, the demonstrated efficacy is due to the product alone; we anticipate even greater efficacy in the presence of ADCC competent bystanders.

A fundamental advantage of stIL15-OPS-γδ system over conventional CAR-T systems is the conferring of enhanced immune activity to non-engineered bystanders, something that chimeric antigen receptors cannot do. To model this more accurately within the engineered product we used computational metrics designed to provide an accurate description of dependence between biological species along a priori defined “edges”. Usually used to describe the strength of interaction between sequential nodes in a signaling pathway, DREMI can also be used to show the dependence between transgene expression and the abundance of downstream markers. To make a meaningful comparison, we focused on markers known to be enhanced by the presence of either transgene. DREMI scores for (transgene)→(marker) were higher for CAR-T than for stIL15-OPS-γδ, suggesting that in CAR-T, signaling is rigidly bound to CAR expression, which is not the case in stIL15-OPS-γδ . To map this behavior across the entire observed network and integrate signal magnitude we plotted co-dependence against signal strength; CAR-T signals enriched in the “strong, dependent”region of the plot, whereas stIL15-OPS-γδ signals mapped to “strong, independent” regions.

The enhancement of non-engineered bystanders was not limited to cells within the original engineered product. Both OPS-γδ and stIL15-OPS-γδ supernatant conferred enhanced antigen-specific cytotoxicity on multiple cell types known to engage opsonized targets. The addition of stIL15 to the construct licensed NK cell engagement which was not achieved by OPS-γδ, reflecting the profound enhancement of NK activity following IL15 stimulus. stIL15-OPS-γδ producing an irrelevant opsonin were still able to receive antigen-specific enhancement if the correct opsonin was provided, demonstrating that the engineering process does not “hijack” ADCC mechanisms and cripple innate γδ T cytotoxicity. Of particular interest was the complete clearance of GD2+ targets by M1 macrophages, suggesting that the cytotoxic capacity of bystander myeloid cells in the microenvironment could be engaged by stIL15-OPS-γδ cells. From a mechanistic perspective, whilst the engagement of opsonized targets by NK cells and M1 macrophages has been well characterized in the past, γδ T cell ADCC remains more ambiguous due to conflicting reports surrounding FcγR expression on γδ T cell subsets*(3, 5, 50)*. By separating staining for the stimulatory CD16a and inhibitory CD16b, we demonstrated a key crossover which occurs during Vδ 2 γδ T cell transduction; whilst expression of CD32 and CD64 remained constant, there was a switch from a predominantly CD18a^-^/CD16b^+^ phenotype, to a predominantly CD16a^+^/CD16b^-^ phenotype, to the extent that the entirety of CD16 expression was CD16a by D6 after transduction. Previous studies of Vδ 2 γδ T ADCC have used high concentrations of opsonin, which masks the dependence on CD16, whereas at opsonin concentrations commensurate with that produced by stIL15-OPS-γδ, CD16^high^ donors have a definite advantage over CD16^low^ donors. Engineering did not appear to have a significant impact on overall CD16 expression, and there was also little impact in terms of Vδ 2 memory phenotype (as determined by CD27/CD45RA expression) compared to unmodified-γδ .

The purpose of an ADCC-engaging T-cell therapy is to remodel the local environment in favor of tumor killing. To do this effectively, the cells must home to the tumor bed. Modelling this in mice is problematic due to the lack of stromal compatibility, though we have already demonstrated in-vivo efficacy against subcutaneous xenografts. Moving to a more translationally relevant system, we evaluated the differential expression of chemokines in normal bone and osteosarcoma to determine those which were enriched in the tumor. Transcriptomic analysis of paired samples from 18 patients revealed enhanced expression of 8 chemokines in the tumor; the receptors for 5 of which were upregulated in stIL5-OPS-γδ . This suggests that engineered cells would have the capacity to seek out and home to areas of bony disease. To evaluate this in an entirely human system we chose a combination of 2D and 3D co-culture methods. Modelling tumour-immune interactions in 3D is an attractive alternative to murine modelling when there is no direct homolog between human and murine immune components. Patient derived osteosarcoma cultures were established in growth-factor stripped Matrigel, with growth factor supplemented media added on top. In order to exert and antitumour response, γδ T added to each well must penetrate and traverse the 3D matrix, homing towards the regions where osteosarcoma has established. This approach allows simultaneous evaluation of signaling and cell-state in effectors and targets within the same system, providing valuable mechanistic insight into the tumour-immune crosstalk that would be much harder to interpret in a murine model. Osteosarcoma suppressed signaling and proliferation in unmodified-γδ, reducing IdU incorporation in particular; stIL15-OPS-γδ resisted this pressure, retaining the enhanced signaling and proliferative markers observed in their target-free state. Presence of OS significantly increased perforin and pSTAT3 abundance, and where GD2 was present, enhanced pAKT levels. Concurrent analysis of the co-cultured osteosarcoma cells demonstrated enhanced levels of perforin and granzyme B within the GD2-PDOS19, and enhanced levels of Granzyme B in GD2+ PDOS25. In both cases, co-culture with stIL15-OPS-γδ led to increased cleavage of PARP and caspase 9 in osteosarcoma, indicative of target apoptosis, a finding not observed in co-culture with unmodified-γδ cells.

Review of the rationale for aminobisphosphonate use in osteosarcoma has traditionally focused on their combination with chemotherapeutic agents, direct anti-sarcoma activity and effects on bone resorption, rather on their role as immunomodulators*(51–53)*. By administering aminobisphosphonates in combination with chemotherapy, any immunological benefit may be masked by bone marrow suppression. In our hands, clinically-achievable doses of zoledronic acid had no effect on osteosarcoma viability. As an adjunct to treatment with γδ T cells, however, zoledronic acid not only enhanced the cytotoxicity of unmodified-γδ T, but also allowed osteosarcoma clearance by stIL15-OPS-γδ . This suggests that a re-evaluation of the value of aminobisphosphonates is warranted - given in the correct combination they may still have a valuable role to play in primary bone tumors such as osteosarcoma.

## MATERIALS AND METHODS

### Experimental design

This study’s primary research objective is proof-of-concept for a novel allogeneic cell-based therapy specifically designed for solid tumour targeting. The therapy comprises Vγ9Vδ 2 γδ T cells engineered to secrete ADCC/ADCP-inducing tumour-targeting opsonins in combination with a cytokine-based mitogen. As this therapy is allogeneic, and can be generated from pre-validated donor material, we designed experiments using γδ T cells sourced from healthy donor blood. We observed a strong correlation between Vδ 2 ADCC efficacy and the amount of CD16 expressed at baseline on the donor’s Vδ 2. In keeping with a proposed “off the shelf” immunotherapeutic, we therefore preselected donors whose Vδ 2 expressed high levels of CD16 for this study. Healthy donor bystander cells were used to demonstrate recruitment of non-engineered immune cells and model potential interactions with the patient’s own immune system. A range of relevant in vitro and in vivo assessments were used to test the feasibility and efficacy of this platform technology, and different techniques including flow cytometry, mass cytometry and ELISA, were employed to interrogate the immune-tumour interactome. To model the tumour, we used both primary tumour lines and isogenic cell lines with engineered GD2 expression. 2D and 3D in vitro experiments were conducted in both experimental and biological replicates until statistical significance was achieved; specific details for the value of n are provided in the figure legends for individual experiments. For our in vivo model, a study was initially conducted with 3 mice per group to inform safety and power calculations, and then repeated with larger group sizes of 6 mice per group. Statistical analyses were performed on complete expanded data sets incorporating both experimental and biological replicates. Investigators were not blinded during experimental setup or sample acquisition, and no outliers were excluded from the datasets presented.

### Cell lines and tissue culture

The following human cell lines were from ATCC: SupT1 (T cell Lymphoblastic Lymphoma), KELLY (Neuroblastoma), Capan-1 (Pancreatic Adenocarcinoma), and Raji (Burkitt’s Lymphoma), HEK293T & HEK293T/17 (human embryonic kidney). SupT1-GD2 were a kind gift from Prof. John Anderson. Culture conditions are described in the supplementary methods. Patient derived osteosarcoma cells designated PDOS16, PDOS19 and PDOS25 were a kind gift from Dr Katia Scotlandi via the ITCC P4 consortium.

### Viral vectors

Polycistronic gene constructs were designed using SnapGene 2.8.3. Gene blocks were synthesized by Twist Bioscience (California, USA) and standard restriction enzyme cloning used to insert them into an RD114 pseudotyped SFG gammaretroviral or pCCL.SFFV lentiviral vector which was either produced in house as previously described*(32)* or supplied by Virocell Biologics (London, UK). Lentiviral vectors were titred on HEK293T cells to determine the number of titratable units per ml.

### γδ T cell expansion and transduction

PBMC isolation is described in the supplementary methods. For specific Vδ 2+γδ T cell expansion, PBMCs were isolated as described above. They were cultured in RPMI 1640 medium supplemented with l-glutamine and 10% FCS (v/v) (Gibco, Massachusetts, USA). Vδ2^+^ γδ T cell expansion was stimulated using 5 μM zoledronic acid (Actavis, New Jersey, USA) and IL-2 (100 IU/ml; Aldesleukin, Novartis, Frimley, UK), which was added to PBMC suspension after PBMC isolation (day 1). Unless otherwise stated, IL-2 was replenished every 2 to 3 days by removing half of the media from the well and replacing with fresh media containing IL-2 (200 IU/ml).

Gammaretroviral transduction of γδ T cells was carried out in RetroNectin (Takara Bio, Tokyo, Japan)–coated 24-well plates, which were preloaded with viral supernatant. T cells (0.5 × 10^6^) suspended in 0.5 ml of T cell medium + IL-2 (400 IU) were combined with 1.5 ml of viral supernatant and centrifuged for 40 min, 1000g at room temperature. γδ T cell gammaretroviral transduction was performed at day 3-5 of culture. Transduction efficiency was determined by flow cytometry on day 5 after transduction. For lentiviral transductions, the vector was added directly to the expansion culture at an MOI of 2-4. Cells were typically harvested on day 12 unless otherwise stated.

### Bystander cell preparation

Untouched NK cells were isolated from frozen and thawed PBMCs using magnetic bead isolation according to the manufacturer’s instructions (Miltenyi Biotec, Bergisch Gladbach, Germany) and used immediately in co-culture assays. Flow cytometry was used to confirm cells were CD3^-^CD56^+^. ADCP competent macrophages were differentiated from peripheral blood monocytes using a previously described method*(54)*. Light microscopy was used to confirm morphology and adherence to plastic. Flow cytometry was used to confirm expression of CD16 and CD64. Neutrophils were differentiated from CD34+ cord blood cells as described and tri-lobed nuclear morphology confirmed using light microscopy *(55)*. Bystander differentiation protocols are described in the supplementary methods.

### *In vitro* cytotoxicity assays

Effector cells were typically co-cultured for 18h with target cells at an E:T ratio of 1:1 unless otherwise stated in the figure legends. Target cells were pre-labelled with Cell-Trace^™^ Violet (Invitrogen) to differentiate between target and effector cells. Flow cytometry was used to count live target cells and/or assess target cell death within gated tumour cells using viability dyes. If relative cell numbers are expressed, then they are normalized to a monoculture of the cells in question, cultured in the same plate and with the same starting number.

To assess bystander cytotoxicity, cell-free culture supernatants from engineered or unmodified Vδ2 were added to co-cultures of targets and allogeneic bystander effector cells (unmodified γδ, NK cells, macrophages or neutrophils). Following co-culture, cells were harvested and analyzed by flow cytometry as above.

### Flow cytometry

Flow cytometric analysis was performed using an LSRII flow cytometer (BD, New Jersey, USA). Data were collected using BD FACSDiva V8.0.1 and analyzed using FlowJo 10.8.1. Compensation was calculated on the basis of OneComp eBeads (Thermo Fisher Scientific) stained with single-color antibodies. Where absolute cell counts were required, Precision Count Beads (Biolegend) were added to tubes prior to sample acquisition. A list of antibodies used is shown in table S1.

### ELISA

The concentration of IL-15 in cell-free culture supernatants was determined using a standard sandwich ELISA according to the manufacturer’s instructions (Biotechne, Minessota, USA). Data points were interpolated from a standard curve ranging from 15.6 – 1,000 pg/ml of recombinant human IL-15 using a sigmoidal 4-parameter logistic model (Prism 9.1.3, GraphPad, California, USA).

### Mass cytometry

All samples from a given donor and stimulation run were barcoded using the TOB*is* barcoding system *(56, 57)*. Up to 20 samples were therefore stained simultaneously in the same tube in a total volume of 300 μl. Cell fixation, permeabilization, and staining were performed as previously described *(32)*. To ensure maximum comparability between samples, all data were acquired using internal metal isotope bead standards (EQ Beads, Fluidigm). A list of antibodies used and their conjugates is shown in Table S2. Mass cytometric acquisition was performed using a Helios mass cytometer (Fluidigm). Approximately 100,000 events were acquired per sample—totalling 2 × 10^6^ events for a full barcoded set.

### CyTOF data postprocessing

Individual time series were normalized to the internal bead standards using the method described in*(58)*, and bead events were removed from the resulting FCS files. In addition, as described in*(59)*, abundance values reported by the mass cytometer were transformed using a scaled arcsinh, with a scaling factor of 5. Debarcoded FCS files were postprocessed using a combination of FlowJo V10.8.1 and bespoke Python scripts. Gating was performed in FlowJo V10.8.1, and gated populations were exported before computational analysis.

Where mass cytometry was used to determine differences in the abundance of species between two samples in a barcoded set, this was achieved using Earth Movers Distance (EMD) calculation. EMD was computed between samples and and donor-matched controls that had undergone the same postprocessing in terms of gating of specific populations. The python module wasserstein_distance, which is a component of scipy.stats, was used to calculate EMD between samples.

DREMI and DREVI analysis DREMI and DREVI are information theory–based methods developed to quantify and visualize relationship between two molecular epitopes*(33)*. Given two proteins epitopes X and Y, and assuming that we are interested in assessing the influence of X on Y, then DREVI visualizes the conditional dependence of Y on X. Where DREVI plots and DREMI scores were calculated, the methodology described in *(32)*, which uses the open source tool simpledremi (Pe’er lab github) was used to derive the scores and generate plots.

### Mouse experiments

All murine studies were approved by the UK home office (PP5675666). Female 6 week-old NOD.Cg-*Prkdc^scid^ Il2rg^tm1Wjl^/SzJ* (NSG) mice were purchased from Charles River (Wilmingtion, MA, USA) and housed in individually ventilated cages with a maximum of 5 mice per cage. SupT1-GD2 were stably transduced with eGFP and firefly luciferase in house using a lentiviral vector and single colony sorting. Mice were challenged with a single subcutaneous injection of 1×10^6^ GFP/firefly luciferase-expressing SUPT1-GD2 cells in 50μls of Matrigel® (Corning, New York, USA) and tumour engraftment confirmed using bioluminescence. 5-7 days later, mice were treated with or without 10×10^6^ non-transduced or transduced Vd2 cells via intravenous injection into the tail vein. Interim bloods were collected 7 days later. Tumour growth was assessed using bioluminescence. Once palpable, tumour size was measured using vernier callipers.

### Differential expression analysis of transcriptomic data

Transcriptomic data on osteosarcoma and normal bone was obtained from publicly available repositories*(41, 60)* Differential expression analysis was performed using DESEq2*(61)*, implemented in Python, using a curated set of chemokines. Data were expressed as Log2 fold change and adjusted p-values corrected for multiple comparison.

### Statistical analyses

All statistical analyses were performed using GraphPad Prism 9.3.0. P values of less than 0.05 were considered statistically significant. *, **, *** and **** were used to indicate p values of <0.05, <0.01, <0.001 & <0.0001, respectively in the figures. Where two groups of values were compared, student’s t-test was used. If more than two groups were compared, one-way ANOVA with Tukey’s multiple comparison correction was used to determine differences between groups. Where other analyses have been used, details are given in the appropriate legend.

## Supporting information

All Supplementary Materials

