## Supplementary material for "OPS-γδ: allogeneic opsonin-secreting γδT cell immunotherapy for solid tumours mediates direct and bystander immunity": All Supplementary Materials

#### List of Supplementary Materials

1. **Supplementary materials and methods**
2. **Supplementary Table 1 – flow cytometric antibodies used**
3. **Supplementary Table 2 – mass cytometric antibodies used**
4. **Supplementary Figures**

### Supplementary Materials & Methods

#### Cell culture conditions

SupT1, Kelly and Raji were maintained in a base medium of RPMI-1640 (Sigma Aldrich, Missouri, USA), whereas Capan-1 were maintained in Isocove's Modified Dulbecco;s Media (IMDM, Sigma Aldrich). HEK293T or HEK293T/17 were used for virus production and were maintained in a base media of Dulbecco's Modified Eagle Medium (Sigma Aldrich). Patient derived osteosarcoma lines were maintained in a base medium of IMDM (Sigma Aldrich).

All base media were supplemented with 2mM L-Glutamine (Sigma-Aldrich) and 10% heat inactivated FBS (v/v) (Thermo Fisher Scientific, Massachusetts, USA).

Where 3D cultures were performed, primary osteosarcoma lines were cultured in growth-factor stripped Matrigel, above which was placed growth-factor supplemented media ABEGNRSW as described in*(75)*. All cell culture including co-cultures was carried out at 37°C with 5% $CO_2$.

#### PBMC preparation

This study was approved by a national ethics committee (West Midlands HRA, 14/WM/1253) and complied with the WMA declaration of Helsinki. Healthy whole blood or leukopaks were collected at the ICH or purchased from Cambridge Bioscience (Cambridge, UK). PBMCs were isolated by density adjusted centrifugation using Lymphopure™ density gradient medium (Biolegend, California, USA). Residual red blood cells and platelets were sequentially removed using ammonium-chloride-potassium lysis solution (Thermo Fisher Scientific) and slow-speed centrifugation, respectively. PBMCs were either used fresh or frozen in a freezing solution containing 10% DMSO as a cryoprotectant. Storage of PBMCs complied with the Human Tissue Act 2004.

#### Bystander cell preparation and identification

Untouched NK cells were isolated from frozen and thawed PBMCs using magnetic bead isolation according to the manufacturer's instructions (Miltenyi Biotec) and used immediately in co-culture assays. Flow cytometry was used to confirm cells were $CD3^{-}CD56^{+}$.

ADCP competent macrophages were differentiated from peripheral blood monocytes. Monocytes were positively isolated using CD14+ microbeads (Miltenyi Biotec) and allowed to adhere to tissue culture plates for 2 hours in sera-free RPMI-1640 at a density of $2\times10^5$ cells/cm$^2$. Sera-free media was then replaced with complete media and the cells cultured at 37°C with 5% $CO_2$ for 10 days. 25ng/ml of recombinant human IFN-g (Biotechne) was added on day 7 to enhance CD16 and CD64 expression and license ADCP function.

Light microscopy was used to confirm morphology and adherence to plastic. Flow cytometry was used to confirm expression of CD16 and CD64.

Neutrophils were differentiated from $CD34^+$ cord blood stem cells. $CD34^+$ stem cells were isolated from cord blood samples obtained from the NHS Blood and Transplant service using $CD34^+$ microbeads (Miltenyi Biotec) and cultured for 3-4 days in X-VIVO 10 plus 1% HSA supplemented with 100 ng/ml stem cell factor, 100 ng/ml human Flt3-ligand, and 100 ng/ml thrombopoietin (all from Peprotech (London, UK)). Cells were then differentiated into granulocytes by culturing for 20 days in Iscove's modified Dulbecco's media supplemented with 20% FBS, 20 ng/ml stem cell factor, 20 ng/ml IL-3 and ng/ml G-CSF 100 (all from Peprotech). Light microscopy was used to confirm tri-lobed nuclear morphology.

**Phenotyping antibody list:**

| Antigen | Fluorochrome conjugate | Clone | Supplier |
|---|---|---|---|
| Vδ2 | APC | REA771 | Miltenyi Biotec |
| CD3 | Vioblue | REA613 | Miltenyi Biotec |
| CD16 | PE-Vio770 | REA423 | Miltenyi Biotec |
| REA control | PE-Vio770 | REA293 | Miltenyi Biotec |
| CD64 | Brilliant Violet™ 605 | 10.1 | Biolegend |
| CD206 | PE | 15-2 | Biolegend |
| CD68 | APC | Y1/82A | Biolegend |
| HLA-DR | Brilliant Violet™ 711 | L243 | Biolegend |
| CD192(CCR2) | PE | K036C2 | Biolegend |
| CD184(CXCR4) | APC-Cy7 | 12G5 | Biolegend |
| CD183(CXCR3) | Brilliant Violet™ 605 | G025H7 | Biolegend |
| CD195(CCR5) | PerCP-Cy5.5 | HM-CCR5 | Biolegend |
| CD186(CXCR6) | PerCP-Cy5.5 | K041E5 | Biolegend |
| CD197(CCR7) | PE | G043H7 | Biolegend |
| CD45RA | Brilliant Violet™ 421 | O323 | Biolegend |
| CD27 | Brilliant Violet™ 711 | HI100 | Biolegend |
| PD-1 | PE | EH12.2H7 | Biolegend |
| TIM-3 | FITC | F38-2E2 | Biolegend |
| LAG-3 | PECy7 | 11C3C65 | Biolegend |
| TIGIT | Brilliant Violet™ 605 | A15153G | Biolegend |
| Anti-IgG | Alexa Fluor® 647 | N/A | Invitrogen |
| Anti-Fc | PE | M1310G05 | Biolegend |

**Mass Cytometry Antibodies**

Osteosarcoma/γδT cell co-culture

| Metal | Antigen / Target | Antibody Clone | Supplier |
|---|---|---|---|
| 116-Cd | GFP | 5F12.4 | eBiosciences |
| 122-Te | TOB*is* Barcode | - | Prof. Mark Nitz |
| 123-Te | TOB*is* Barcode | - | Prof. Mark Nitz |
| 124-Te | TOB*is* Barcode | - | Prof. Mark Nitz |
| 125-Te | TOB*is* Barcode | - | Prof. Mark Nitz |

| | | | |
|---|---|---|---|
| 126-Te | TOB*is* Barcode | - | Prof. Mark Nitz |
| 127-I | IdU | - | Fluidigm |
| 128-Te | TOB*is* Barcode | - | Prof. Mark Nitz |
| 130-Te | TOB*is* Barcode | - | Prof. Mark Nitz |
| 142-Nd | Cleaved-Caspase 3 [D175] | D3E9 | CST |
| 143-Nd | CD45 | HI30 | BioLegend |
| 145-Nd | Phospho-ZAP70 [Y319] | 17a | BD Biosciences |
| 149-Sm | CD274 (PD-L1) | 29E.2A3 | BioLegend |
| 150-Nd | Phospho-RB [S807/811] | J112-906 | BD Biosciences |
| 152-Sm | Phospho-AKT [T308] | J1-223.371 | BD Biosciences |
| 155-Gd | CD3 | UCHT1 | BioLegend |
| 156-Gd | Phospho-NF-κB p65 [S529] | K10-895.12.50 | BD Biosciences |
| 158-Gd | CD27 | L128 | BD Biosciences |
| 167-Er | Phospho-ERK1/2 [T202/Y204] | 20A | BD Biosciences |
| 169-Tm | Phospho-STAT3 [Y705] | 4/P-Stat3 | BD Biosciences |
| 170-Er | TCR γδ | B1 | BioLegend |
| 171-Yb | CD44 | IM7 | BioLegend |
| 172-Yb | CD16 | B73.1 | BioLegend |
| 173-Yb | Granzyme B | QA16A02 | BioLegend |
| 174-Yb | Cleaved-PARP [D214] | D64E10 | CST |
| 175-Lu | Perforin | DG9 | BioLegend |
| 191-Ir | DNA | - | Fluidigm |
| 193-Ir | DNA | - | Fluidigm |

stIL15-γδ phenotyping

| Metal | Antigen / Target | Antibody Clone | Supplier |
|---|---|---|---|
| 116-Cd | GFP | 5F12.4 | eBiosciences |
| 122-Te | TOB*is* Barcode | - | Prof. Mark Nitz |
| 123-Te | TOB*is* Barcode | - | Prof. Mark Nitz |
| 124-Te | TOB*is* Barcode | - | Prof. Mark Nitz |
| 125-Te | TOB*is* Barcode | - | Prof. Mark Nitz |
| 126-Te | TOB*is* Barcode | - | Prof. Mark Nitz |
| 127-I | IdU | - | Fluidigm |
| 128-Te | TOB*is* Barcode | - | Prof. Mark Nitz |
| 130-Te | TOB*is* Barcode | - | Prof. Mark Nitz |
| 141-Pr | CD253 (TRAIL) | RIK-2 | BioLegend |
| 142-Nd | Cleaved-Caspase 3 [D175] | D3E9 | CST |
| 143-Nd | CD45 | HI30 | BioLegend |
| 144-Nd | Phospho-STAT5 [Y694] | 47/Stat5 | BD Biosciences |
| 145-Nd | Phospho-ZAP70 [Y319] | 17a | BD Biosciences |
| 146-Nd | Phospho-SLP76 [Y128] | J141-668.36.58 | BD Biosciences |
| 147-Sm | Phospho-BTK [Y551] | 24a/BTK | BD Biosciences |
| 148-Nd | Phospho-SRC [Y418] | SC1T2M3 | Thermo |
| 149-Sm | CD274 (PD-L1) | 29E.2A3 | BioLegend |
| 150-Nd | Phospho-RB [S807/811] | J112-906 | BD Biosciences |
| 152-Sm | Phospho-AKT [T308] | J1-223.371 | BD Biosciences |
| 153-Eu | TIGIT (VSTM3) | A15153G | BioLegend |
| 155-Gd | CD3 | UCHT1 | BioLegend |
| 156-Gd | Phospho-NF-κB p65 [S529] | K10-895.12.50 | BD Biosciences |
| 157-Gd | CD366 (Tim-3) | F38-2E2 | BioLegend |

| | | | |
|---|---|---|---|
| 158-Gd | CD27 | L128 | BD Biosciences |
| 159-Tb | CD69 | FN50 | FN50 |
| 161-Dy | CD279 (PD-1) | EH12.2H7 | BioLegend |
| 163-Dy | CD226 (DNAM-1) | 11A8 | BioLegend |
| 166-Er | CD337 (NKp30) | P30-15 | BioLegend |
| 167-Er | Phospho-ERK1/2 [T202/Y204] | 20A | BD Biosciences |
| 168-Er | CD178 (Fas-L) | NOK-1 | BioLegend |
| 169-Tm | Phospho-STAT3 [Y705] | 4/P-Stat3 | BD Biosciences |
| 170-Er | TCR γδ | B1 | BioLegend |
| 171-Yb | CD223 (LAG-3) | 7H2C65 | BioLegend |
| 172-Yb | CD16 | B73.1 | BioLegend |
| 173-Yb | Granzyme B | QA16A02 | BioLegend |
| 174-Yb | Cleaved-PARP [D214] | D64E10 | CST |
| 175-Lu | Perforin | DG9 | BioLegend |
| 191-Ir | DNA | - | Fluidigm |
| 193-Ir | DNA | - | Fluidigm |
| 196-Pt | TOB*is* Barcode | - | BuyIsotope |
| 198-Pt | TOB*is* Barcode | - | Fluidigm |
| 209-Bi | CD336 (NKp44) | P44-8 | BioLegend |

### Supplementary Figures

A

| *Antigen* | *scFv* | *Tumour* | *Reference* |
|---|---|---|---|
| GD2 | 14G2a | Glioma, neuroblastoma, sarcoma | (19) |
| CEACAM5 | SM3EL | Breast, lung, ovary, prostate, pancreatic | (20,21) |
| CD20 | 10H9.06 | B cell malignancies | (22) |

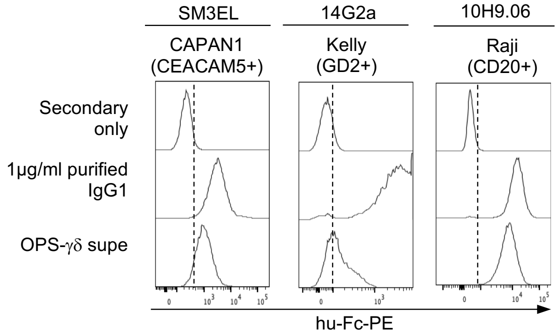


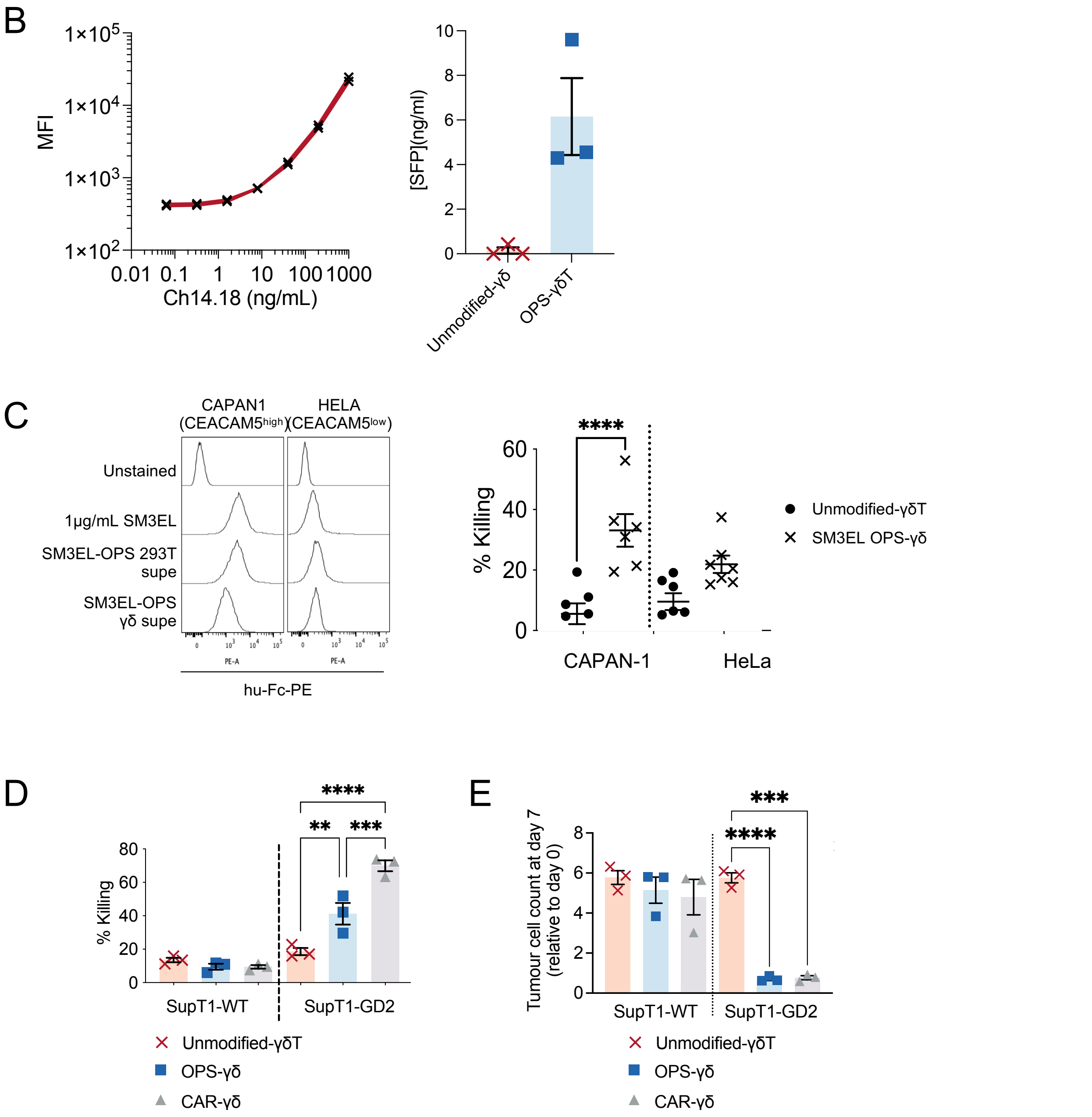

**Supplementary Figure 1**

**(A)** To target a range of malignancies, three different SFP constructs were created, specific for either GD2, CEACAM5 or CD20, and designated 14G2a, SM3EL and CD20, respectively. Flow cytometry of various antigen-positive tumour cells exposed to culture supernatant from either unmodified-γδT, 14G2a OPS-γδ, SM3EL OPS-γδ or 10H9.06 OPS-γδ. Fluorochrome-conjugated anti-Fc or anti-IgG were used to detect SFP binding. 1ug/ml purified whole IgG1 against the relevant antigen was used as a positive control. Representative plots from 3-5 donors are shown. (**B)** Detection and quantification of 14G2a opsonin in supernatant from 14G2a-γδ cultured at D12 of culturewas achieved using flow cytometry. SupT1-GD2 stained with reducing concentrations of Ch14.18, which comprises the same binder in whole antibody format followed by anti-human Fc secondary staining were used to generate a standard curve (left) which was used to quantify scFv-Fc fusion protein in supernatant (right). **(C**) Staining of CAPAN-1 and HELA cells with SM3EL (anti-CEACAM5) scFv-Fc fusion protein binders produced in Vδ2 γδT cells or HEK293T, detected using anti-human Fc secondary antibody. Cytotoxicity of SM3EL OPS-γδ or unmodified γδ against CEACAM5$^{hi}$ CAPAN1 or CEACAM5$^{lo}$ HELA (n=5-6 across 5 donors) **(D)** Percentage killing of SupT1-WT and SupT1-GD2 cells by unmodified-γδ or 14G2a OPS-γδ (n = 13 across 8 donors) or 14G2a CAR-γδ (G, n=3 donors) in an overnight co-culture at an E:T of 1:1. (**E)** Long term growth assay for SupT1-WT and SupT1-GD2 cells in co-culture with either unmodified-γδ, OPS-γδ or CAR-γδ. (n=3 donors).

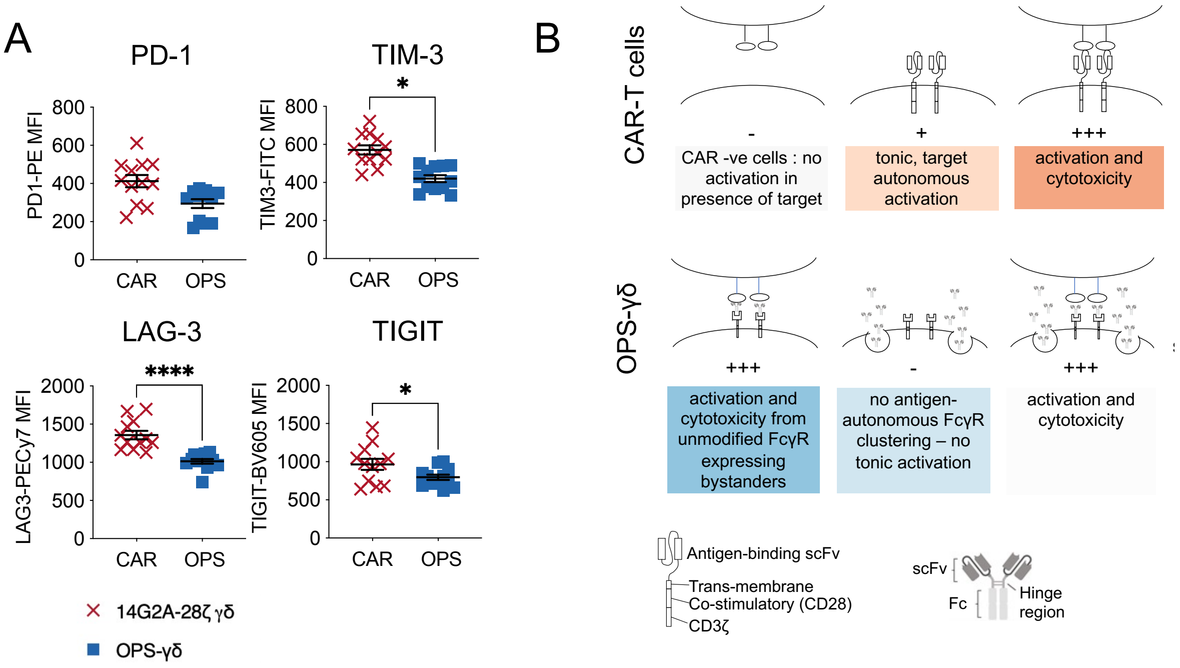


**Supplementary Figure 2**

**(A)** MFI of PD-1, TIM-3, LAG-3 and TIGIT on live $CD3^{+}V\delta2^{+}$ cells within 14G2a OPS-γδ and CAR-γδ on day 5 after transduction (n=12 across 3 donors). All graphs show individual data points with mean ± SEM, comparison by unpaired t-test. **(B)** Schematic comparing and contrasting OPS-γδ with CAR-γδ.

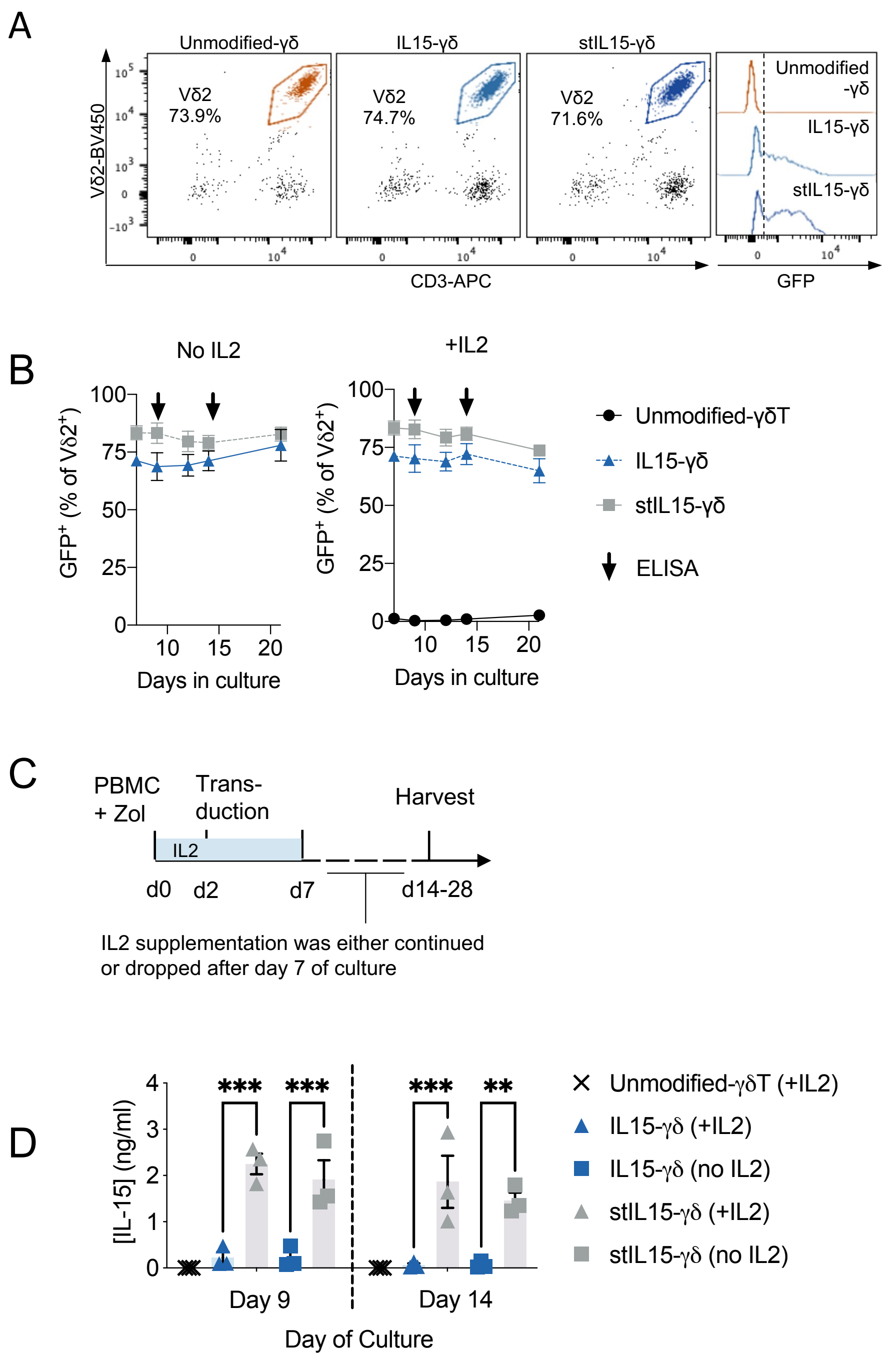


**Supplementary figure 3**

**(A)** Representative flow cytometry data from 6 donors showing γδT purity and transduction efficiency.using either GFP-2A-IL15 or GFP-2A-stIL15 **(B)** Percentage GFP expression within live $V\delta2^{+}CD3^{+}$ cells from unmodified-γδT, IL15-γδ and stIL15-γδ between day 7 and 21 of culture as measured by flow cytometry. Data points are means±SEM for n=3 donors **(C)** Schematic depicting the experiment used to test the expansion of IL15-γδ and stIL15-γδ in the absence of exogenous cytokine support. **(D)** Concentration of IL15 in culture supernatants from unmodified-γδT, IL15-γδ and stIL15-γδ at day 9 and 14 of culture as determined by ELISA. “+IL2” and “no IL2” in parenthesis indicates whether IL2 supplementation was dropped at day 7 of culture or not. Individual data points and means±SEM are shown for n=3 donors

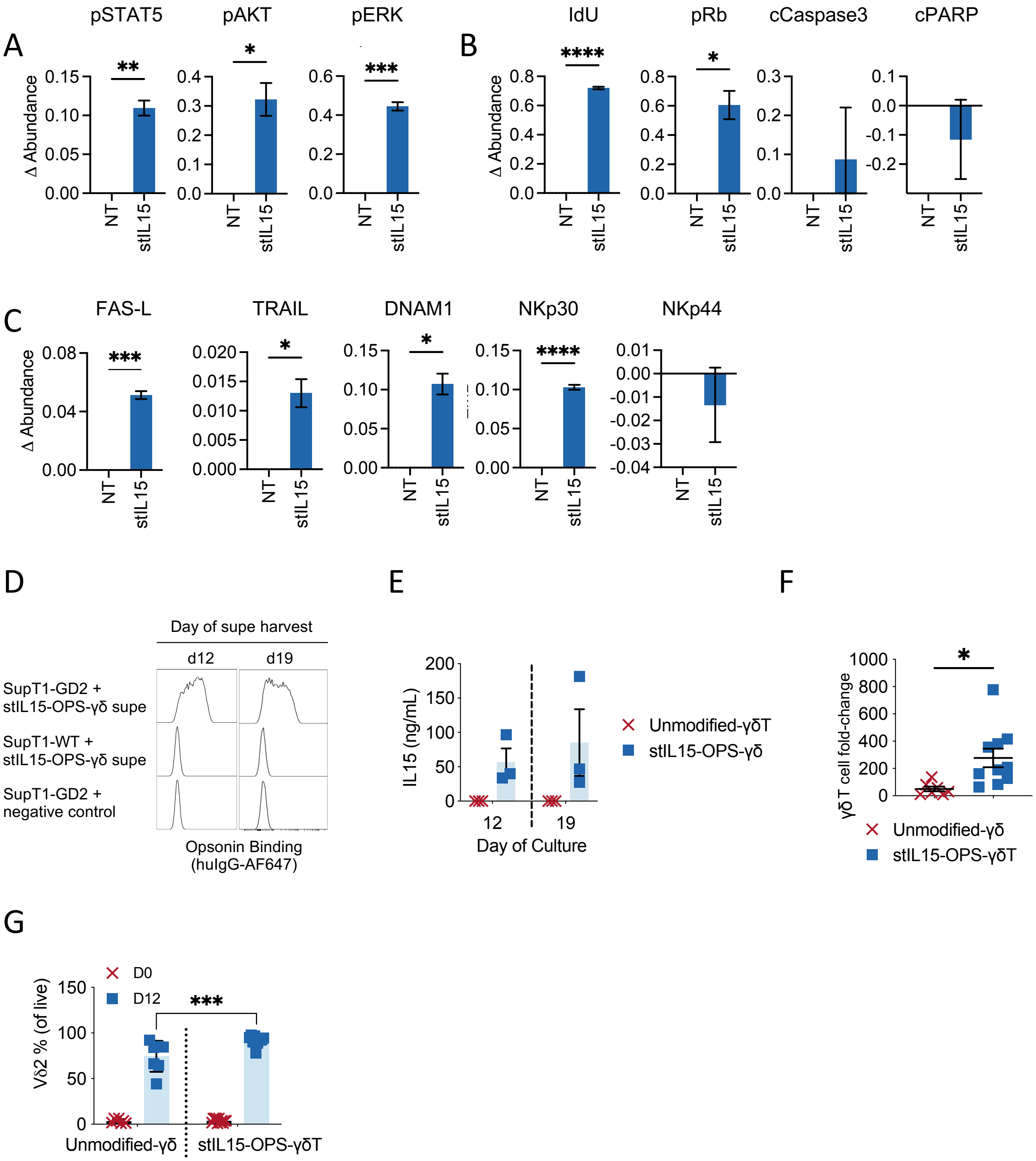


**Supplementary figure 4**

A-C: Expression of signaling, cell state and phenotypic markers was assessed in unmodified-□□T and stIL15-γδ on day 12 of expansion using mass cytometry. EMD was used to express the difference between stIL15-γδ and unmodified γδ, so differences are normalized to unmodified γδT cells as the baseline. Means±SEM for n=5 are shown. **(A)** Differences in abundance of signaling markers in stIL15-γδ compared to unmodified γδ. **(B)** Differences in abundance of markers of proliferation and apoptosis in stIL15-γδ compared to unmodified γδ. **(C)** Differences in abundance of innate cytotoxicity markers in stIL15-γδ compared to unmodified γδ. **(D)** Flow cytometry of SupT1-WT or SupT1-GD2 cells exposed to either day 12 or day 19 culture supernatant from stIL15-OPS-γδ, followed by fluorochrome-conjugated anti-IgG.

Representative plots from 6 donors are shown. **(E)** The concentration of IL15 in day 12 or day 19 culture supernatants from unmodified-γδT or stIL15-OPS-γδ as determined by ELISA. Individual data points and means±SEM are shown for n=3 donors **(F)** Fold change in the number of Vd2+CD3+ cells within unmodified γδ and stIL15-OPS-γδ between day 0 and day 12 of expansion as determined by flow cytometry and cell counts. Individual data points and means±SEM are shown for n=7-10 over 5 donors. **(G)** Percentage Vd2+CD3+ cells of gated live cells within unmodified γδ and stil15-γδ at day 0 and 12 of expansion as determined by flow cytometry. Individual data points and means±SEM are shown for n=7-11 over 5 donors.

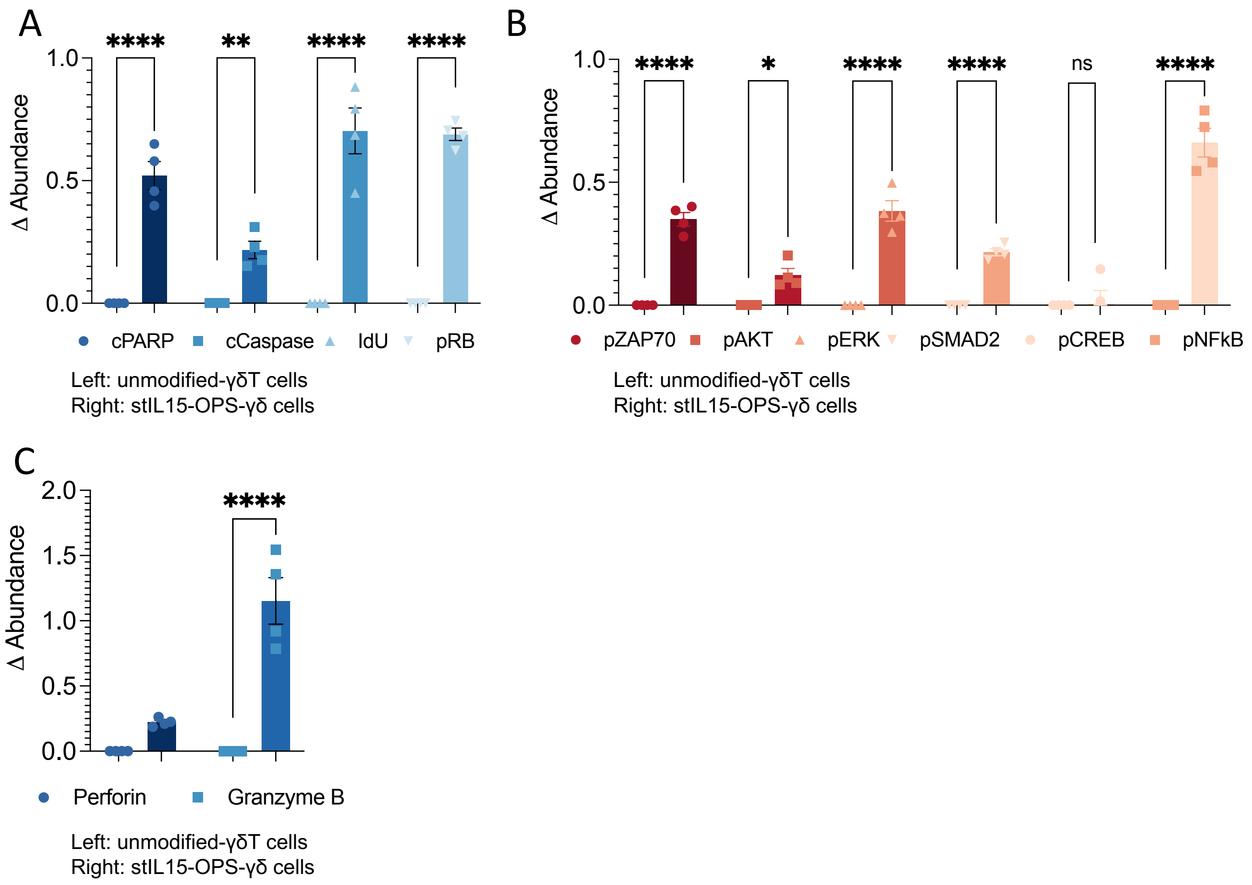


**Supplementary figure 5**

Expression of key markers of proliferation & apoptosis **(A),** activation & ITAM signaling **(B)** and cytotoxic function **(C)** was assessed in unmodified γδ and stIL-15-γδ on day 12 of expansion using mass cytometry. Differences are with reference to unmodified-γδ as the baseline, and graphs show mean EMD±SEM of n=4 across 2 donors.

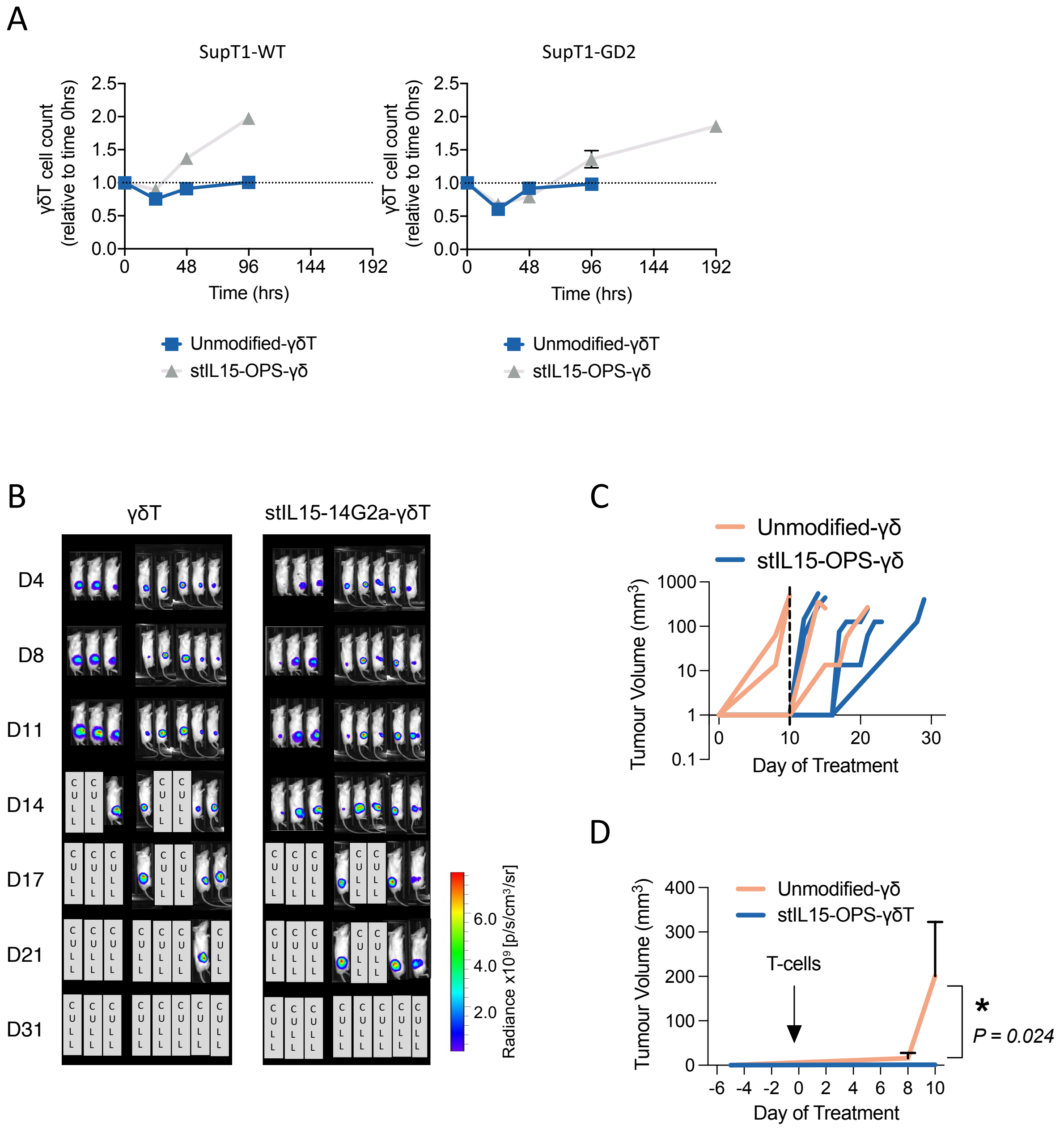


**Supplementary figure 6**

**(A)** Vδ2+ cell counts during the challenge and re-challenge assays illustrated in (C), comparing stIL15-OPS-γδ or unmodified γδT cells (mean ± SEM of n=6 across 2 donors). **(B)** Bioluminescence images of NSG mice bearing subcutaneous, luciferase expressing SupT1-GD2 tumors following intravenous injection of $1x10^7$ unmodified γδ or stIL15-14G2a-γδ. **(C)** Subcutaneous SupT1-GD2 tumour volumes in individual NSG mice for the entire duration of the experiment **(D)** Mean volumes±SEM up to D10 following intravenous injection of $1x10^7$ unmodified γδ or stIL15-14G2a-γδ are also shown.

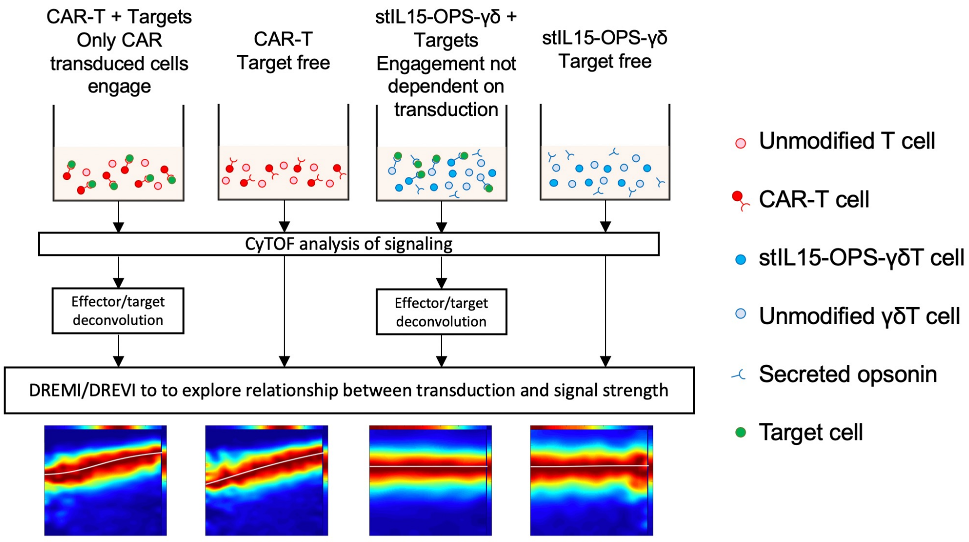


**Supplemental Figure 7**

Culture and analysis setup used to determine the dependence between transgene expression and markers of activation at single-cell resolution.

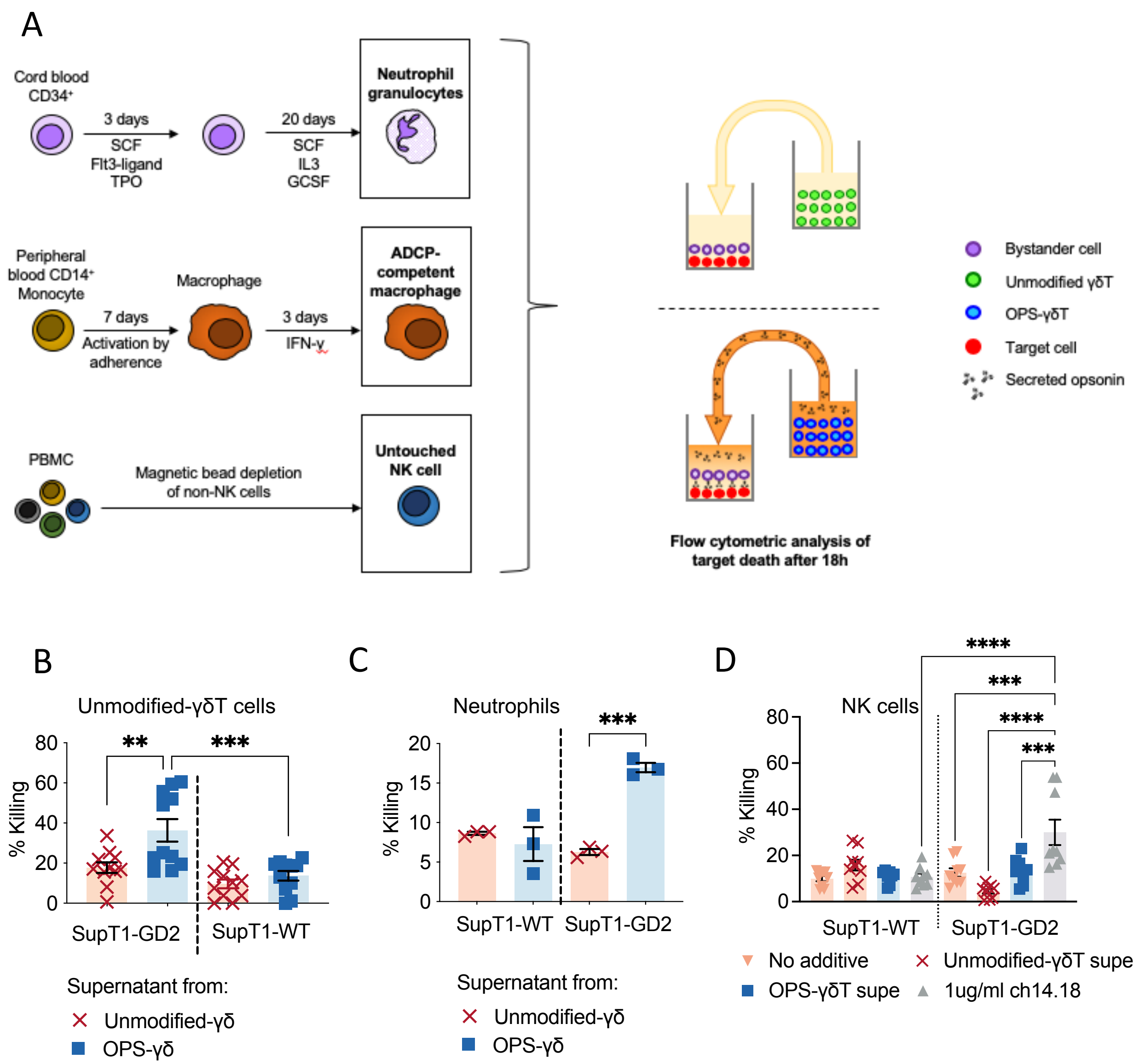


**Supplementary Figure 8**

**(A)** Schematic of setup for generating neutrophil, macrophage and NK cell bystanders and then demonstrating their engagement by products contained in supernatant from OPS-γδ or stIL15-OPS-γδ. Flow cytometry was used to measure the effect of supernatant from unmodified γδT or OPS-γδ on percentage killing of SupT1-WT and SupT1-GD2 cells co-cultured with various effectors including unmodified expanded γδT (**B,** n=9 across 5 donors), (**C**) Stem cell-derived neutrophils (mean ± SEM of n=3 supernatants from 3 donors) and (**D**) resting NK cells (n=9 supernatants from 3 donors or n=9 for ch14.18 control).

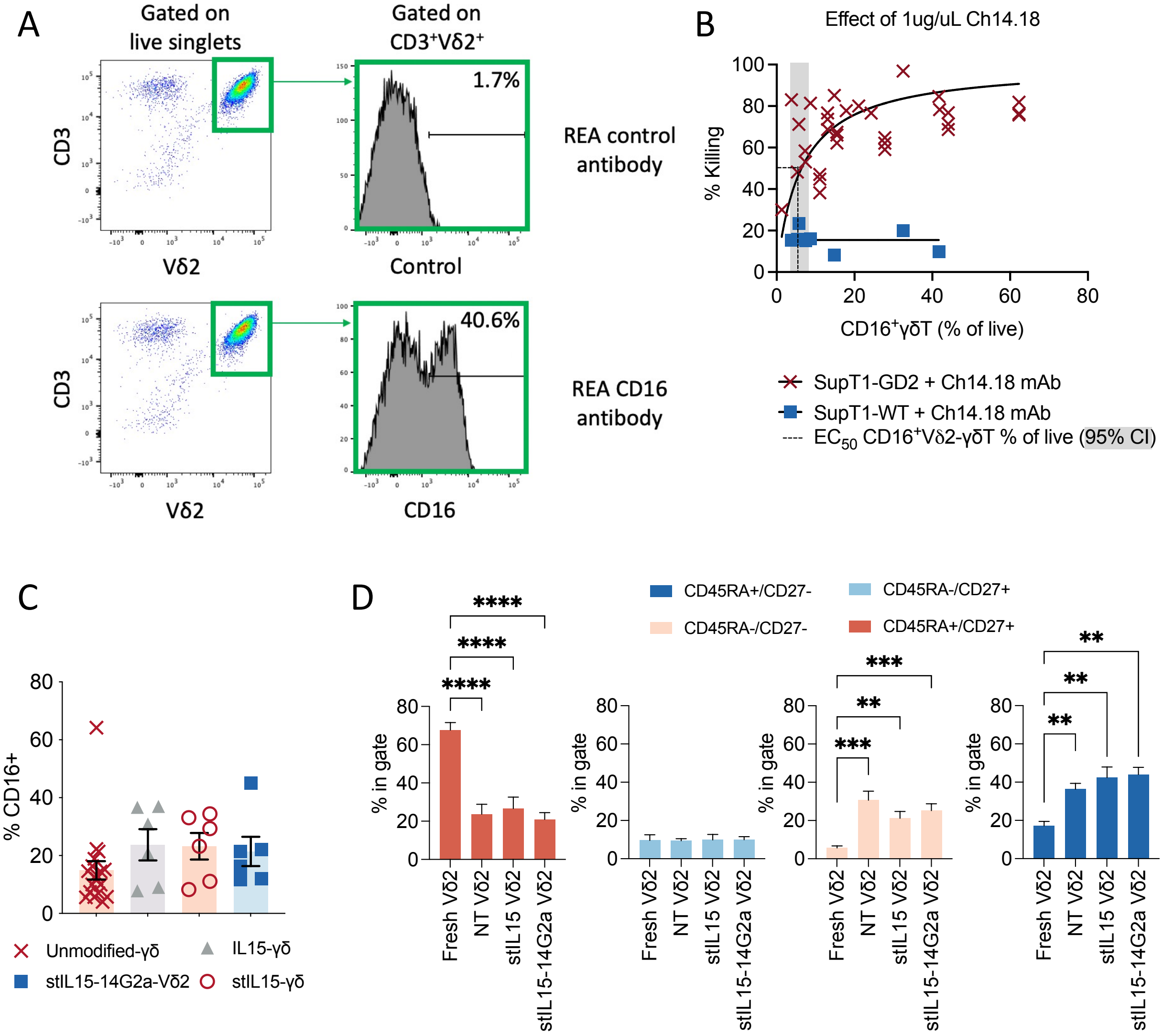


**Supplemental Figure 9**

**(A)** Example gating for determination of % CD16 (FcγRIII) expression on Vδ2 γδT cells. **(B)** Killing of SupT1-WT or SupT1-GD2 by unmodified γδ in the presence of 1μg/ml Ch14.18, plotted against levels of FcγRIII (CD16) on the corresponding donor's Vδ2 after expansion. Least squares curve fitting was used to determine the $EC^{50}$ for CD16 expression, n=32 across 19 donors for SupT1-GD2 and n=8 across 8 donors for SupT1-WT. **(C)** Percentage of FcγRIII expression on γδ cells within unmodified-γδT, IL15-γδ, stIL15-γδ and stIL15-OPS-γδ on day 12-14 of expansion as determined by flow cytometry. Individual data points and means±SEM for n=6 donors **(D)** Flow cytometry was used to assess CD45 and CD27 expression on live Vd2+CD3+ cells within fresh PBMC or unmodified gd, stIL15-gd and stIL15-OPS-γδ at day 12 of expansion. The percentage of $CD45^{+}CD27^{-}$, $CD45^{-}CD27^{-}$, $CD45^{-}CD27^{+}$ and $CD45^{+}CD27^{+}$ within gated Vd2+CD3+ cells is shown. Bars represent means±SEM are shown for n=12 across 6 donors.

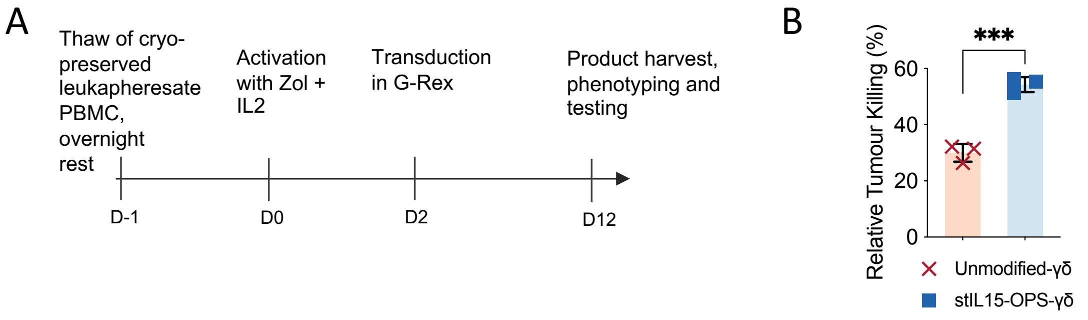


**Supplementary Figure 10**

**(A)** Process for expanding and transducing Vδ2 γδT cells from frozen PBMC stocks using scalable, GMP compatible G-Rex vessels. **(B)** Cytotoxicity of G-Rex expanded (as in (A)) stIL15-14G2a-γδ or unmodified γδ against SupT1-GD2

A PDOS25

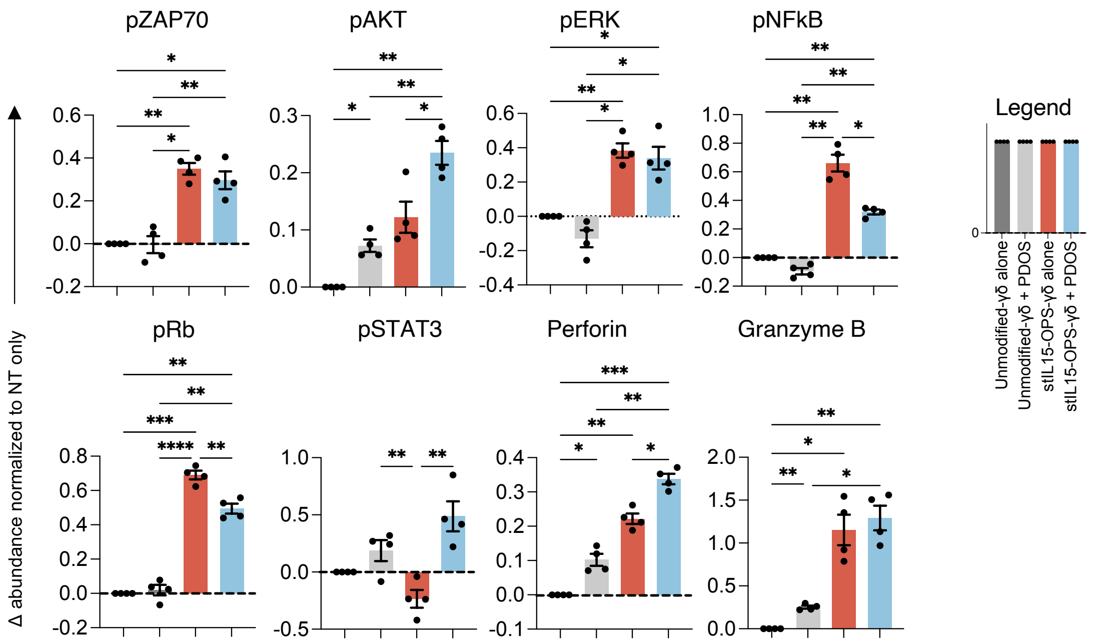


B PDOS19

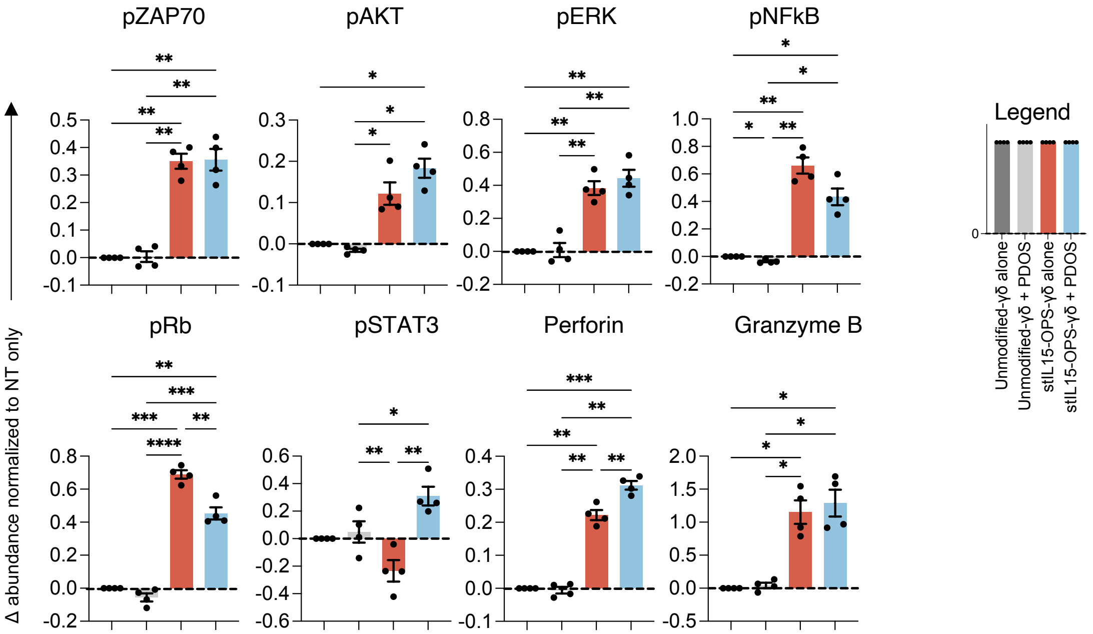


**Supplementary Figure 11**

**(A)** GD2 expression patterns of 3 primary osteosarcoma lines PDOS16, PDOS19 and PDOS25. Differences in the expression of activation signal and cytotoxicity markers in unmodified γδ and stIL15-14G2a-γδ in the presence or absence of GD2 expressing PDOS25 (**B**) or GD2 negative PDOS19 **(C)** as determined by mass cytometric analysis of γδT cells in 3D culture. Differences were computed using Earth Movers Distance, with donor matched unmodified γδ as the baseline. Data shown are means±SEM of n=4 across 2 donors.
